# From Benchmark to Bench: Can Agents Survive Real-World Drug Discovery?

**DOI:** 10.64898/2026.09.30.755603

**Authors:** Pierre Llompart, Levent Guner, Helen Lai, Alessandro Tibo, Yijie Xu

## Abstract

Agentic systems increasingly coordinate molecular-design tools, but it is unclear which layer of the stack limits outcomes on real projects. We developed MAGI, an open modular agent that authors objectives, launches and monitors optimization, interprets structure–activity relationships, and revises its strategy accordingly. MAGI generates molecules either directly through the LLM or by delegating to REINVENT 4, with scoring services interchangeable behind a common contract. We tested it across nine retrospective lead-optimization campaigns from three pharmaceutical companies, replayed under fixed temporal cutoffs. Both routes produced valid structures: LLM proposals stayed closer to local chemistry and reached comparable or higher primary activity in fewer operations, whereas REINVENT explored broader chemical space. Whether a campaign met its objective depended on the predictive models, not on the generation route: attainment followed model accuracy on the chemistry proposed, dropping once that chemistry moved outside the model’s applicability domain. Separately, a blinded evaluation asked whether the MAGI’s output could pass as expert work: chemists were not able to discriminate agentic proposals from held-out compounds, and judged the SAR reasoning broadly plausible yet incomplete. Together, these results position MAGI as a coordination layer pluggable into existing computational chemistry workflows. The ceiling on real projects, however, remains currently set by scorer applicability rather than by tool orchestration.

## 1 Introduction

Lead optimization advances a compound toward a candidate by balancing potency, safety and developability across repeated design–make–test–analyze cycles, a major contributor to drug-development cost [1]. Generative methods such as REINVENT 4 optimize molecules against expert-defined objectives [2, 3], but their scoring configurations require revision as project evidence and structure–activity relationships evolve.

Agentic systems promise to manage this process by planning, invoking tools, and adapting in closed loops [4–6]. Chemistry agents and orchestrators have recently proliferated in literature [7–22], yet most evaluations test tool execution or generic molecular benchmarks rather than scientific performance under real project constraints. Strong benchmark performance can coexist with elementary reasoning failures and unreliable confidence [23–25], while language-model molecular generation alone guarantees neither molecular validity nor controlled multiparameter optimization [26]. An agentic campaign is steered by its scoring services, which act as oracles: every decision the agent makes is expressed through them. Their applicability therefore bounds what the agent can achieve. Systematic bias shifts the numerical scale, range compression suppresses real differences, and poor rank correlation makes candidate ordering unreliable. Across the nine campaigns reported here, attainment tracks the applicability of models and success from generation route.

MAGI is an open, modular agent that authors and adapts project objectives, launches and monitors optimization campaigns, and interprets emerging SAR to redirect them. Molecular generation occurs either directly through the LLM or through agent-controlled REINVENT 4, with both routes using a common prediction and structure-evaluation stack. We evaluate MAGI on four GSK series [27–30], four Sanofi projects [31], together with one undisclosed AstraZeneca campaign, all retrospectively replayed under fixed temporal cutoffs. Our contributions are:

- An auditable modular agent architecture in which project objectives are executable, versioned, and shared across LLM and REINVENT generation routes.
- A first-time evaluation on nine industry campaigns from three pharmaceutical companies reporting validity, objective attainment, workload, and limitations.
- An evidence that objective attainment on drug discovery projects is bound by the reliability of predictive models rather than by agent coordination.

## 2 Related works

REINVENT 4 is a widely used production framework for goal-directed molecular generation and optimization [2]. It maximizes an expert-defined scoring function but does not author or revise that objective, making it a natural generator within an agentic workflow. LLM molecular design systems differ in where generation and critique occur. MT-Mol generates SMILES through specialized analysis, design, and verification agents [18], whereas CIDD combines an LLM medicinal-chemistry critic with pocket-conditioned 3D generators [13]. DrugR uses explicit pharmacological reasoning [16]; MolClaw combines REINVENT 4, Boltz-2 and interaction profiling [20]; Mozi uses governed agents for molecular optimization [11]; and ChatInvent integrates REINVENT and synthesis planning in a production-facing interface [10]. Validation on temporally held-out project chemistry remains rare. RetroDMTA provides four time-resolved industrial programs [31], but its original benchmark prioritizes known future compounds. MAGI instead exposes only pre-cutoff context during generation and uses future compounds for retrospective comparison.

## 3 Method

### 3.1 System

MAGI separates agent orchestration, molecular generation, and evaluation. Claude Sonnet 4.6 accesses REINVENT 4 [2], global ADMET models, temporally restricted project models [31], Boltz-2 [32], PoseBusters [33], and an interaction profiler through a common REST contract exposed by the Model Context Protocol [34]. Project objectives are represented as versioned therapeutic target profiles (TTPs) that compile into executable REINVENT scoring configurations. The MCP server exposes 24 tools covering data access, TTP management, scoring, campaign control, and result retrieval. Services, target constructs, and the architecture are documented in Table S1, Table S4, and Figure S1a.

Optimization jobs execute asynchronously and are monitored by the gateway. Independently containerized services exchange structures, configurations, and results through shared storage. An external REINVENT component batches generated SMILES to these services, while a dependency-aware wrapper passes Boltz-2 complexes to PoseBusters and the interaction profiler. Structural evidence can therefore contribute directly to optimization rather than only post hoc ranking.

### 3.2 Objective and configuration space

Each campaign is governed by a versioned, machine-readable therapeutic target profile. Each objective specifies its scoring service, optimization direction, target value or acceptable interval, desirability transform, and weight; fixed guardrails are evaluated separately as pass/fail constraints. Endpoint predictions are transformed to [0, 1] desirabilities and combined using a weighted Derringer–Suich geometric mean [35]. The complete formulation is given in Section S2.3. The same TTP evaluates LLM proposals and compiles the REINVENT scoring configuration; every agent-authored revision is versioned, and missing values remain unevaluable. Project-specific objectives and guardrails are reported in Table S6.

## 4 Experimental setup

### Datasets

MAGI is evaluated on nine lead-optimization tasks: four published GSK BET BD2 series [27–30], four Sanofi RetroDMTA projects—FXA, MMP8, PPAR*δ* and RENIN [31]—and one undisclosed AstraZeneca campaign. Each task shows the agent one starter and four measured analogues and withholds every remaining compounds. The primary endpoints are kinetic solubility for GSK-S1, membrane permeability for GSK-S2 and log *D*_7.4_ for GSK-S3 and GSK-S4; log *D*_7.4_ for AstraZeneca; p*K*_i_ for FXA, pIC_50_ for MMP8 and RENIN, and pEC_50_ for PPAR*δ*, with solubility and log *D*_7.4_ as additional FXA objectives. Sanofi project models see only pre-cutoff compounds, so the future cohort remains held out at both generation and scoring. Endpoint definitions, cohort sizes and splits are reported in Tables S2 and S3.

### Protocol

Every campaign ran for five iterative optimization rounds. Each round supplies the project context and the current objective, generates and evaluates molecules, and ends with an agent-authored shortlist of approximately five compounds carried into the next round. On the REINVENT route the agent additionally creates or updates a TTP, launches and monitors optimization, and retrieves the generated pool from which it selects those compounds. Figure S1b summarizes the protocol, and Tables S7 and S8 report pool sizes, shortlist construction, cofolding coverage, and generation parameters.

### Metrics

Molecular quality is summarized by validity, cumulative uniqueness, novelty to the shown set, similarity to held-out chemistry, intra-round diversity and synthetic accessibility [36, 37], where diversity is one minus the mean pairwise ECFP4 Tanimoto similarity. Cofolding is summarized by Boltz-2 ligand iPTM and interaction counts. Operational load is the number of observed generation, prediction, cofolding, validation, monitoring, retrieval and inspection calls. Project- and route-level results are reported in Figure 1 and Figures S4–S16.

**Figure 1:**
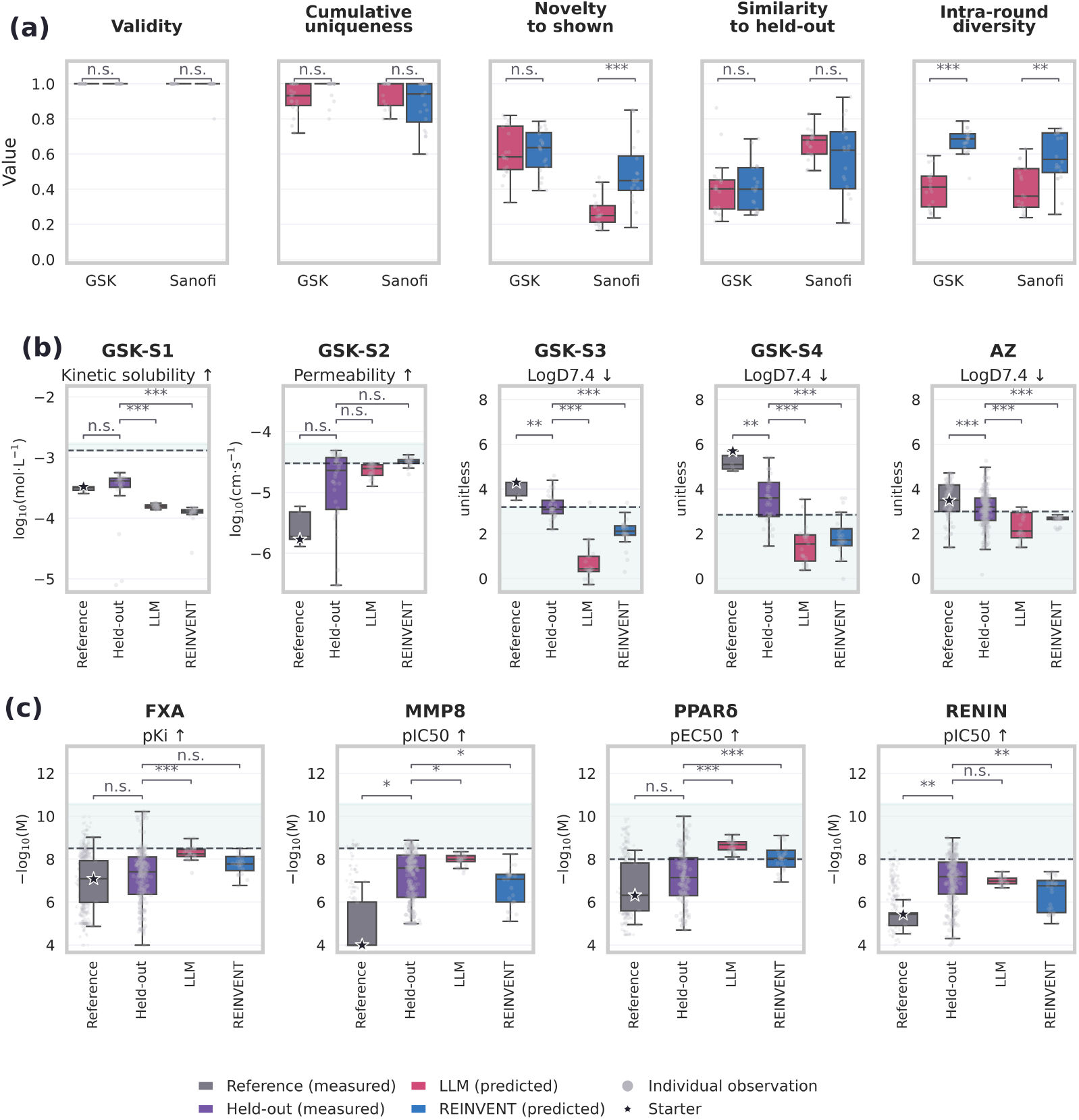
Molecular quality and objective attainment across Drug Discovery campaigns. (a) Project-round validity, cumulative uniqueness, novelty, held-out similarity, and intra-round diversity for LLM (red) and REINVENT (blue). (b) GSK and AstraZeneca primary-endpoint distributions. (c) Sanofi project-level primary-endpoint distributions. Pale-green shading indicates the objective-satisfying region. Pale points denote pre-cutoff Reference compounds. Stars mark the selected starter compound. Brackets show Holm-adjusted two-sided Mann–Whitney U tests, adjusted p-value (*p*) significance (∗ *p* < 0.05, ∗∗ *p* < 0.01, ∗ ∗ ∗ *p* < 0.001; n.s., not significant).

## 5 Results

### 5.1 Molecular quality and chemical-space exploration

We first assess the validity and uniqueness of proposals from each generative route. Figure 1a compares round-level validity, cumulative uniqueness, novelty to shown chemistry, similarity to held-out chemistry and intra-round diversity across the GSK and Sanofi projects. Route differences depended on dataset and objective. GSK proposals were more novel overall, and within Sanofi it was REINVENT that departed furthest from the shown chemistry; conversely, Sanofi proposals resembled the held-out compounds more closely than GSK ones did. REINVENT achieved greater intra-round diversity in both organizations, supporting broader exploration.

### 5.2 Primary-endpoint and objective-attainment evaluation

Chemical validity alone does not establish whether a campaign advances its project objective. We therefore compare each route’s proposals against two experimental references on the primary endpoint: the analogues shown to the agent, and the held-out compounds it never saw (Figure 1b for GSK and AstraZeneca, Figure 1c for Sanofi).

Three of the four GSK series and the AstraZeneca campaign met their objective. GSK-S1 was the exception: its kinetic-solubility target sits outside the training range (Table S5). During the experiment, the agent autonomously promoted thermodynamic solubility as a proxy, followed by an improvement in proposals (Figure S12). On the Sanofi campaigns, both routes matched the held-out compounds on predicted primary activity, yet only FXA and PPAR*δ* cleared the project threshold. The LLM reached the objective more often than REINVENT, reproducing the quality these programmes historically reached without surpassing it. In case where a campaign fell short, the cause coincided with a model operating outside its applicablity range: GSK-S1 failed because its scorer was extrapolating, while RENIN ligand-based model showed weak held-out correlation (Figure S3). Supporting analyses report round-wise diversity (Figure S4), synthetic accessibility and chemical-space (Figures S9–S11), models validation (Figure S3), trajectories (Figures S12–S16) and molecules (Figures S5–S8).

### 5.3 Operational behaviour and medicinal chemistry evaluation

The routes produced similar profiles but set different operational demands. Figure 2 summarizes operations and adaptation events. REINVENT required four-to-nine times more operations than the LLM (Figure S17), arising from launch, monitoring, recovery, and retrieval. These counts exclude the calls issued inside the reinforcement-learning loop: each REINVENT campaign explored 100,000 more structures than the LLM, increasing the number of tool calls to four more orders of magnitude. MAGI’s adaptations extended beyond syntax repair: it revised objectives, search parameters, and molecular strategy in response to convergence, scorer disagreement, and emerging SAR (Section S2.6). In a blinded evaluation involving four chemists and 28 questions, participants were unable to distinguish AI-generated from held-out molecules, showed no preference between design routes, and found the agent’s SAR reasoning plausible but incomplete (Section S6.3, Figure S18).

**Figure 2:**
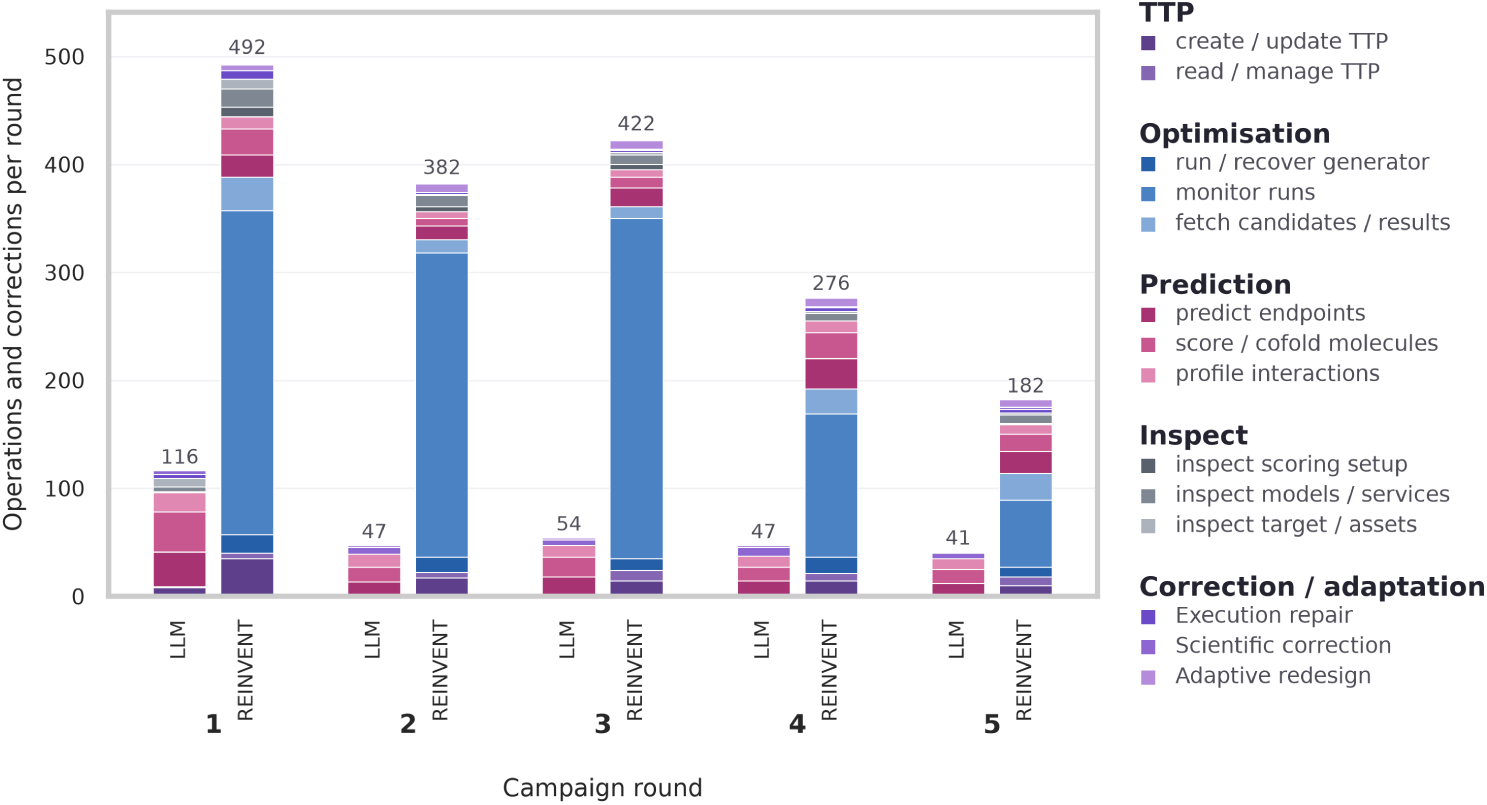
Observable agent operations and correction/adaptation events. Counts are shown by category, campaign round, and generation route; counts exclude scorer and model calls executed internally within REINVENT’s reinforcement-learning loop.

## 6 Discussion

The two generative routes expose an exploration–exploitation trade-off. LLM generation acted as a focused local optimizer around the shown chemistry, whereas agent-controlled REINVENT broadened diversity and novelty when a project needed to leave its starting chemotype. Neither route consistently dominated model-space attainment, indicating that generator breadth cannot compensate for a biased, range-compressed, or weakly ranking scorer. Structure-based services shifted proposal rankings in a target-dependent manner: reranking by Boltz-2 affinity changed the top-5 shortlist substantially (Figures S2 and S3d). The blinded evaluation highlighted that chemists could not separate agent proposals from measured compounds nor that they a preference for held-out over AI compounds. MAGI’s contribution lies above the orchestration, in selecting which tool to invoke and revising the objective as evidence accumulates [18, 21].

## 7 Limitations

Proposal endpoints are predictions, not measurements. Each condition used one backend, one trajectory, and no static-objective control, so the agent’s own contribution to attainment is not isolated. Temporal splits restrict train/test overlap but cannot exclude pretraining exposure.

## 8 Conclusion

We presented MAGI, a modular reasoning layer that versions project objectives, coordinates established generators and predictors, monitors runs, and adapts execution as evidence accumulates. Across nine industrial campaigns, LLM generation provided local optimization with fewer operations, while REINVENT broadened exploration at higher operational cost. Neither route consistently won on objective attainment as temporal controls show this depended on the scoring models: campaigns succeeded when predictors were reliable and failed otherwise. Real project data surfaced this limit in a way benchmarks would not, from weak model performance on industrial data to autonomous adaptation by the agent through proxy endpoints. Overall, agentic design survives contact with real projects, but two critical ceilings remain: the quality of the agent’s SAR reasoning, and the applicability of scoring services in an industrial context.

## Supplementary Information

### S1 Extended related work

Agentic systems for chemistry began with tool-augmented assistants: ChemCrow coupled a language model with chemistry tools [4], Coscientist further leveraged wet-lab automation to drive experiments [5], and multi-agent architectures refined hypotheses [6]. Drug-discovery orchestrators followed, with pipeline assembly from natural language [7, 8], tool selection by tree search [9], production deployment around REINVENT [10], governed hierarchies [11], parameterized memory [12], a critic over a structure-based generator [13], modular task execution [14] and agentic affinity prediction [15]. Agents aimed specifically at lead optimization add explicit-reasoning optimization [16], trajectory-aware tool planning [17], role-specialized multi-agent optimization [18], and a full-process system on the Model Context Protocol [19, 34]. MolClaw is the closest published stack to ours [20–22].

### S2 Agent

#### S2.1 System

The gateway agent runs a bounded tool-use loop (at most 100 tool rounds per agent turn) and writes observable model messages, tool requests, and tool results to a structured JSONL transcript; a background monitor polls job health and injects explicit alerts. The therapeutic target profile (TTP) is a schema-validated, versioned JSON object. Each native or external objective identifies its scoring service, optimization direction, target value or acceptable interval; desirability transforms and, when applicable, weight. Dependencies and aggregation rules define the composite score, while fixed guardrails and SMARTS-based structural constraints are evaluated separately as pass/fail criteria. Each campaign starts with TTP version 1 before the first round. The primary endpoint, data split or temporal cutoff, and fixed guardrails remain unchanged throughout the campaign. Secondary objectives, weights, or search settings may change only through a recorded TTP or configuration update, which increments the stored version. A mapper topologically orders objective dependencies and compiles the TTP into an executable REINVENT scorer configuration [2]. The same TTP semantics are used to evaluate molecules proposed directly by the LLM and molecules generated by REINVENT, ensuring that both routes are compared under the same objectives and guardrails. Each service exposes a POST /score endpoint used by the TTP and by REINVENT scoring, and, where implemented, a POST /predict endpoint that the agent can call directly for analysis. Boltz-2 returns cofolding, affinity, confidence, and structure artifacts [32]; PoseBusters consumes the corresponding upstream complex [33]; the general ADMET and RetroDMTA services return endpoint predictions and uncertainty metadata [31]. Tool exceptions are returned to the agent as errors. External REST scoring failures are logged and represented as NaN unless an explicit failure score is configured. Missing service outputs remain unevaluable and are not counted as objective attainment; a language-model estimate is never substituted for a missing service value. The architecture separates decisions from evidence generation: the gateway chooses a route and manages a versioned objective, while molecular generation, prediction and structural checks remain independently replaceable as services.

### S2.2 Services usage

Both generation routes access the same molecular-evaluation services and author a TTP to score the molecules they produce. The LLM route asks the reasoning backend to propose a shortlist directly; the REINVENT route additionally optimizes the TTP, monitors the job, retrieves the generated pool, and selects the final shortlist from it.

Which services a campaign uses depends on the organization. GSK and AstraZeneca campaigns used the general ADMET stack and excluded cofolding, whereas Sanofi campaigns used the target-specific project model and required Boltz-2/PoseBusters structural evaluation. Boltz-2 produced a complex and an affinity prediction, and PoseBusters consumed that exact complex for geometry evaluation. Every service in Table S1 was reachable either inside the executable TTP, through the common POST/score contract, or directly by the agent for analysis; the general ADMET and project-specific services support both modes. Table S1 reports the deployed versions, outputs, and quality controls. Prediction requests were batch-oriented and preserved input order.

**Figure S1:**
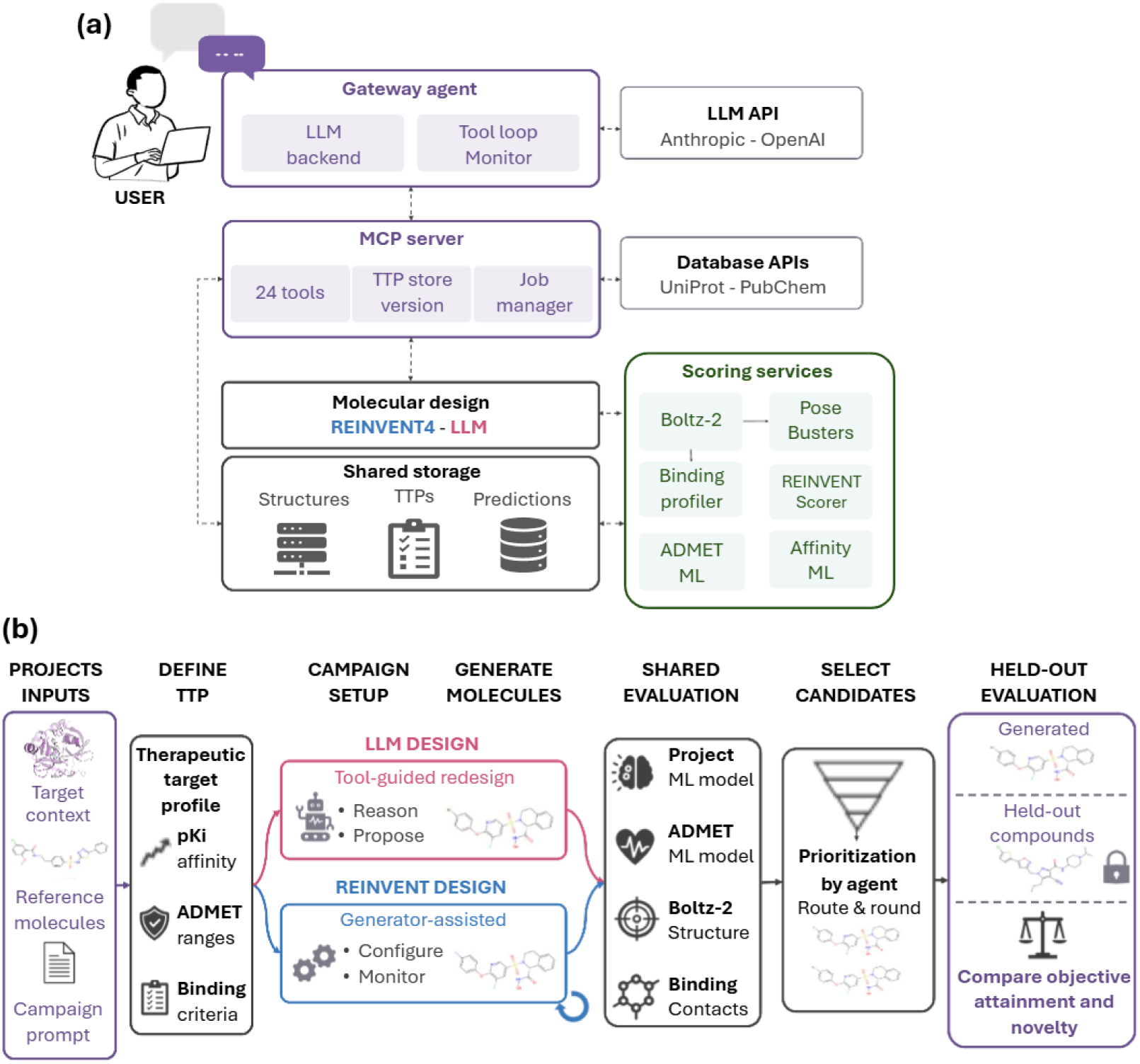
MAGI service interaction, MCP interface, and experimental protocol. **(a) System architecture.** The gateway agent connects the LLM reasoning backend to a monitored tool loop. Through the MCP server, it manages versioned therapeutic target profiles, optimization jobs, REINVENT 4, external data sources, and independently deployable scoring services. Shared storage transfers structures, configurations, and results between services without placing large artifacts in the language-model context. A TTP defines the scoring pipeline; molecules proposed by the LLM or generated by REINVENT are evaluated in batches using native molecular properties, Boltz-2 cofolding, pose and interaction analysis, global ADMET models and project-specific RetroDMTA models. Their component outputs are aggregated and returned to the generation loop to guide the next iteration. The MCP server exposes 24 tools across six groups: service discovery, data retrieval, TTP management, scoring and prediction, optimization control, and result retrieval. **(b) Controlled campaign protocol.** Project context, reference molecules, and objectives are encoded in a versioned TTP. LLM and REINVENT generate candidates through distinct routes but use the same activity, ADMET, Boltz-2, and interaction-scoring services. Ranked evidence guides subsequent rounds, after which final candidates are compared with measured held-out compounds never shown to the agents.

**Table S1:**
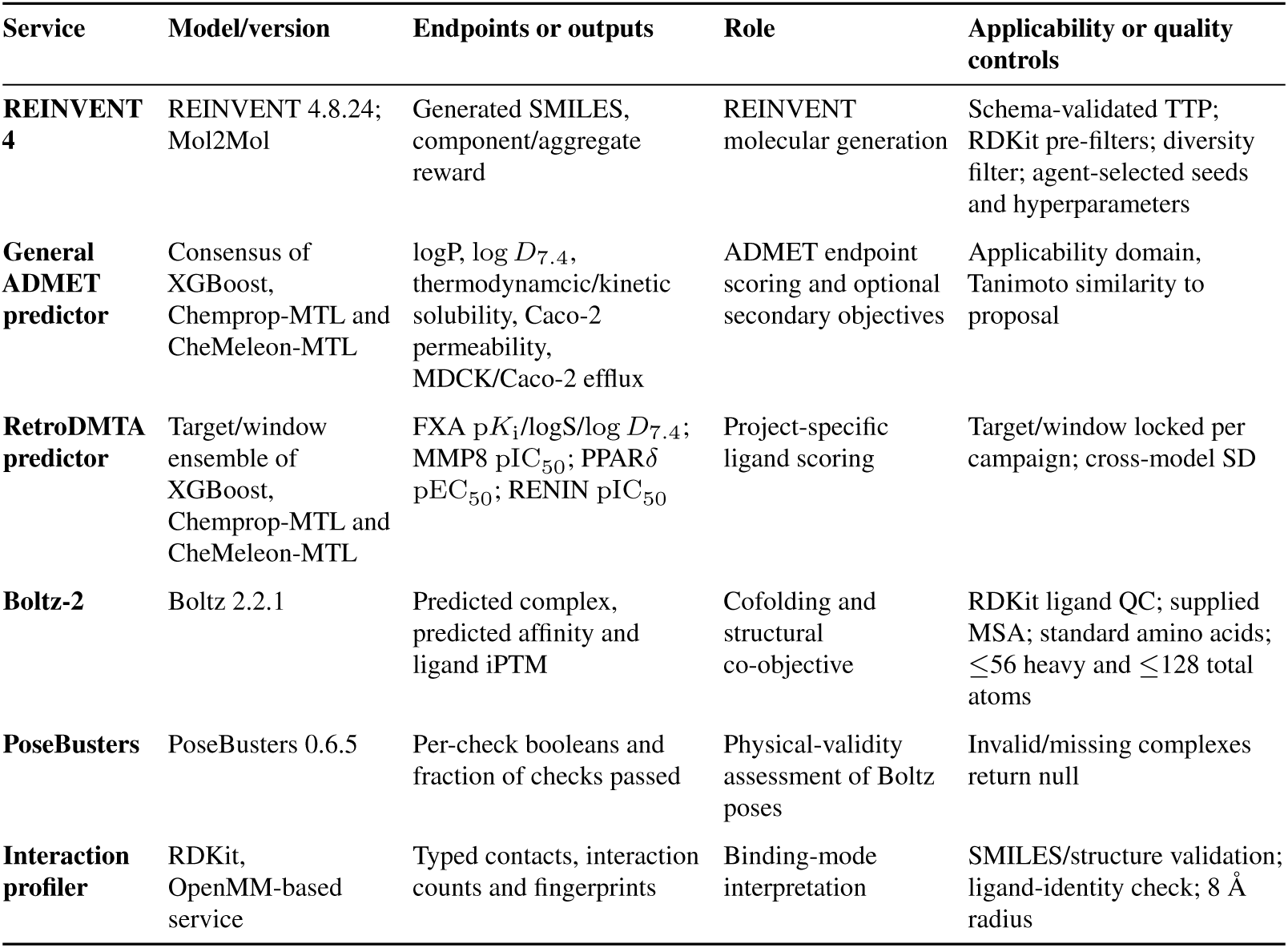
Agent-accessible services, model versions, outputs, roles, and applicability or quality controls.(MTL: Multi-Task Learning)

### S2.3 TTP scoring semantics

For objective *i*, the TTP records a scoring service, optimization direction, target value or acceptable interval, desirability transform *T_i_*, and weight *w_i_*. For molecule *m*, the service prediction *x_i_*(*m*) is transformed to

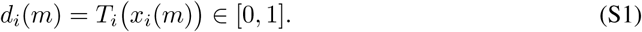

Endpoint desirabilities are combined using the weighted Derringer–Suich geometric mean:

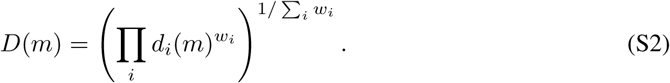

Here, *D*(*m*) is the overall desirability and 1 *− D*(*m*) the distance to the profile. The geometric mean prevents strong performance on one endpoint from compensating for complete failure on another [35]. Complete TTP attainment is reported separately and requires every mandatory objective and fixed guardrail to pass. Missing service values remain unevaluable and are not treated as attainment.

### S2.4 Scoring contract

External scorers use an index-aligned REST contract: POST/score gets a SMILES list and returns scores, metadata, and artifacts. Agent-facing POST /predict endpoints provide analysis without requiring a TTP. The MCP scoring layer formats each molecule with total and component scores, raw and transformed values, failure reasons, warnings, TTP identity/version, and available structure paths. Pipeline dependencies are passed by artifact identity, so PoseBusters consumes the exact Boltz-2 complex associated with each molecule. Network or HTTP failures in the external scorer are logged and produce NaN by default; an explicit native-only fallback is labelled as such.

### S2.5 System prompt

The agent system prompt defines the agent as a medicinal-chemistry orchestration layer, requires tool-grounded molecular claims, and specifies TTP authoring, monitoring, selection and terminal reporting. Per-round task prompts provide project chemistry, objectives, guardrails, and route-specific constraints. The configured backend was Claude Sonnet 4.6; provider-side sampling defaults were used, and no random seed was fixed for the reasoning model. We analyse only observable messages, tool requests, tool results, and configuration payloads.

### S2.6 Agent adaptation, error recovery, and TTP provenance

For every campaign, MAGI records each observable model message, tool request, tool result, TTP version, and executable scoring configuration. Scientific adaptations are evidence-motivated changes to objectives, weights, transformations, search settings, or design strategy and are recorded separately from execution of repairs. For example, if candidates repeatedly fail a solubility guardrail and the agent consequently increases the solubility weight, the guardrail failures are the preceding evidence, and the weight change is the scientific adaptation. A change is classified as a scientific adaptation only when the transcript explicitly links preceding campaign evidence to the subsequent change. Syntax repairs, routine initialization, status polling, and repeated serialization are classified separately as execution work. An operational error is an observable failed or invalid tool interaction, including service unavailability, malformed input, schema-validation failure, job failure, missing artifact, timeout, or failed results retrieval. A recovery is counted when a subsequent observable agent action resolves the error, and the intended operation completes successfully. Recovery is classified as autonomous when no human intervention or manual configuration change occurs between the error and successful completion. Repeated status polling without an error is not counted as recovery.

### S2.7 Compute requirements

Experiments ran in containerized services on nodes equipped with AMD EPYC 9575F CPUs and AMD Instinct MI355X accelerators under ROCm 7.0. REINVENT and Boltz-2 used separate GPUs; structure standardization, PoseBusters and tabular prediction were CPU-parallelized. Tables S7 and S8 report generation workloads and Figure S17 reports input-token volume versus observable operations.

## S3 Datasets

### S3.1 Global ADMET data

Raw experimental records were obtained from OCHEM [38] and ChEMBL [39]. Assay metadata was harmonized before combining sources. log *D*_7.4_ records included chromatographic log *D*_7.4_ measurements. Agreement between project-specific log *D*_7.4_ and global ADMET log *D*_7.4_ is verified in Figure S3a, with the log *D*_7.4_ model demonstrating high accuracy over the GSK project data. Solubility records were restricted to aqueous-buffer measurements at pH 7.4 and separated into thermodynamic aqueous solubility and kinetic solubility measured after dilution of a DMSO stock into buffer. Kinetic-solubility records included measurements obtained after dilution of a DMSO stock into aqueous buffer, including CLND-reported (ChemiLuminescent Nitrogen Detection) measurements when they satisfied the same protocol and unit-curation criteria. Caco-2 permeability retained inhibitor-free apical-to-basolateral (A*→*B) measurements. Efflux measurements were also required to be inhibitor-free and were retained as separate Caco-2 and MDR1–MDCK endpoints. Transport assays used compound concentrations below 100 µM.

After the structure, unit and replicate curation described in Section S3.5, the multitask dataset contained 156,797 unique molecules with one or more labels across seven endpoints: logP, log *D*_7.4_, aqueous thermodynamic solubility, kinetic solubility, Caco-2 permeability, MDR1–MDCK efflux ratio and Caco-2 efflux ratio. The sparse multitask dataset contains 156,797 unique molecules and 186,120 datapoints.

**Table S2:** Global ADMET endpoint definitions, curated datapoint counts, and retained value ranges.

| Endpoint | Conditions | Datapoints | Minimum | Maximum |
| --- | --- | --- | --- | --- |
| LogP | – | 56,140 | –3.0 | 7.0 |
| $\log D_{7.4}$ | PBS Buffer pH 7.4<br>Chromatography | 33,628 | –3.0 | 7.0 |
| Thermodynamic solubility | PBS Buffer $7.4 \pm 0.5$<br>Temperature $25 \pm 5^\circ\text{C}$<br>No excipients or cosolvents | 32,078 | –8.0 | 1.0 |
| Kinetic solubility | Temperature $25 \pm 5^\circ\text{C}$ | 48,116 | –6.5 | –3.7 |
| Permeability Caco-2 | No engineered cell lines<br>No P-gp inhibitors<br>Below 100 $\mu\text{M}$ concentration<br>Below 20 cell passages<br>Apical to basolateral transport | 9,600 | –8.0 | –4.0 |
| Efflux ratio Caco-2 | No engineered cell lines<br>No P-gp inhibitors<br>Below 100 $\mu\text{M}$ concentration<br>Below 20 cell passages | 2,007 | –0.5 | 2.0 |
| Efflux ratio<br>MDCK-MDR1 | No engineered cell lines<br>No P-gp inhibitors<br>Below 100 $\mu\text{M}$ concentration<br>Below 20 cell passages | 4,551 | –0.5 | 2.9 |

### S3.2 BET BD2 ADMET series

The four series were extracted from published GSK lead-optimization studies targeting the second bromodomains of the BET-family proteins BRD2, BRD3, BRD4 and BRDT: the GSK046 series (GSK-S1; [27]), template-hopping series (GSK-S2; [29]), N-methylpyridine-2-carboxamide series (GSK-S3; [28]) and DNA-encoded-library-derived GSK040 series (GSK-S4; [30]). Structures and measurements extracted from the articles and supporting information were processed using the common standardization protocol in Section S3.5.

GSK-S1 uses CLND kinetic solubility, converted from mass concentration to mol/L using the standardized neutral-parent molecular weight, and reported as log_10_(mol/L). This assay conforms to the kinetic-solubility endpoint represented in the general ADMET training set: solubility measured after dilution of a DMSO stock into an aqueous buffer.

GSK-S2 uses artificial-membrane apparent permeability, converted from nm/s to log_10_(cm/s). The GSK-S2 source endpoint is membrane apparent permeability, whereas the deployed general model predicts Caco-2 apparent permeability. The latter was used as a directional permeability surrogate rather than as an assay-identical reconstruction of the published endpoint.

GSK-S3 and GSK-S4 use log *D*_7.4_, retained as the reported dimensionless log_10_ distribution coefficient. The general ADMET model was trained on the corresponding pH-7.4 log *D*_7.4_ endpoint and included chromatographic log *D*_7.4_ records meeting the same endpoint definition.

Each task presents one starter and four measured analogues to the agent and withholds the remaining published structures. Repeated source measurements were aggregated by the median; missing endpoint values were excluded without imputation.

### S3.3 AstraZeneca campaign

The AstraZeneca campaign targets log *D*_7.4_ on an undisclosed internal programme. The compound set, the target, and the therapeutic area are confidential and are not released; only aggregate distributions and the campaign objective are reported here. Reference and held-out cohorts were supplied as measured log *D*_7.4_, aqueous solubility and affinity values without structures, and were processed under the same unit conventions as Section S3.5. Cohort sizes and ranges are reported in Table S3; the three endpoints are not measured for all endpoints. The campaign used the general ADMET stack, and excluded cofolding.

### S3.4 RetroDMTA projects

The four Sanofi datasets represent projects targeting activated coagulation factor X (FXa), matrix metalloproteinase-8 (MMP-8), peroxisome proliferator-activated receptor delta (PPAR*δ*), and RENIN [31]. Fixed temporal cutoffs produced the following pre-cutoff and future cohorts: FXa, 2004-07-28 (803 and 212; 1,015 total); MMP-8, 2003-04-17 (204 and 226; 430 total); PPAR*δ*, 2006-05-08 (355 and 143; 498 total); and RENIN, 2007-02-14 (147 and 241; 388 total).

The primary endpoints are p*K*_i_ for FXa, pIC_50_ for MMP-8 and RENIN, and pEC_50_ for PPAR*δ*, where p*X* = *−* log_10_(*X* in mol/L). FXa additionally contains aqueous thermodynamic solubility at pH 7.4, supplied in µM and converted as log *S* = log_10_(mol/L), and log *D*_7.4_ measured at pH 7.4. Standardization preserves CPD_ID and temporal membership. Each CPD_ID occurs once, so no repeat aggregation is applied; missing endpoint values are excluded without imputation.

**Table S3:** Experimental endpoint composition by dataset and cohort. Ranges and medians exclude missing values; no missing endpoint is imputed.

| Project | Endpoint | Set | N | Min | Max | Unit |
| --- | --- | --- | --- | --- | --- | --- |
| GSK-S1 | Kinetic solubility | Reference | 5 | -3.593 | -3.481 | $\log_{10}(\text{mol/L})$ |
| GSK-S1 | Kinetic solubility | Held out | 31 | -5.104 | -2.882 | $\log_{10}(\text{mol/L})$ |
| GSK-S2 | Permeability | Reference | 5 | -5.886 | -5.229 | $\log_{10}(\text{cm/s})$ |
| GSK-S2 | Permeability | Held out | 46 | -6.523 | -4.31 | $\log_{10}(\text{cm/s})$ |
| GSK-S3 | $\log D_{7.4}$ | Reference | 5 | 3.5 | 4.3 | $\log_{10} D$ |
| GSK-S3 | $\log D_{7.4}$ | Held out | 38 | 2.2 | 4.6 | $\log_{10} D$ |
| GSK-S4 | $\log D_{7.4}$ | Reference | 5 | 4.8 | 5.7 | $\log_{10} D$ |
| GSK-S4 | $\log D_{7.4}$ | Held out | 38 | 1.45 | 5.4 | $\log_{10} D$ |
| AZ | $\log D_{7.4}$ | Reference | 89 | 1.4 | 4.717 | $\log_{10} D$ |
| AZ | $\log D_{7.4}$ | Held out | 262 | 0.1692 | 5.256 | $\log_{10} D$ |
| AZ | Aqueous solubility | Reference | 88 | -7.0 | -3.0 | $\log_{10}(\text{mol/L})$ |
| AZ | Aqueous solubility | Held out | 264 | -7.0 | -3.0 | $\log_{10}(\text{mol/L})$ |
| AZ | Affinity | Reference | 102 | 4.0 | 8.489 | p-scale |
| AZ | Affinity | Held out | 843 | 4.0 | 8.497 | p-scale |
| FXA | $pK_i$ | Reference | 803 | 4.0 | 10.0 | p-scale |
| FXA | $pK_i$ | Held out | 212 | 4.0 | 10.2218 | p-scale |
| FXA | Aqueous solubility | Reference | 803 | -7.0 | 0.3324 | $\log_{10}(\text{mol/L})$ |
| FXA | Aqueous solubility | Held out | 212 | -7.0 | 0.4031 | $\log_{10}(\text{mol/L})$ |
| FXA | $\log D_{7.4}$ | Reference | 803 | -3.989 | 6.665 | $\log_{10} D$ |
| FXA | $\log D_{7.4}$ | Held out | 212 | -3.62 | 5.77 | $\log_{10} D$ |
| MMP8 | $pIC_{50}$ | Reference | 204 | 4.0 | 8.8861 | p-scale |
| MMP8 | $pIC_{50}$ | Held out | 226 | 5.0 | 8.8814 | p-scale |
| PPARδ | $pEC_{50}$ | Reference | 355 | 4.4815 | 9.8765 | p-scale |
| PPARδ | $pEC_{50}$ | Held out | 143 | 4.699 | 10.0 | p-scale |
| RENIN | $pIC_{50}$ | Reference | 147 | 4.0 | 8.3979 | p-scale |
| RENIN | $pIC_{50}$ | Held out | 241 | 4.0 | 9.0 | p-scale |

### S3.5 Data processing and sample flow

Structures were parsed and sanitized with RDKit. Salts, cosolvents and surfactants were removed, and the largest organic fragment was retained, selected by heavy atom count and then molecular weight. Supplied stereochemistry was preserved during aromatic reperception and functional-group normalization. Structures were converted to the RDKit canonical tautomer, assigned their dominant ionization state at pH 7.4 and represented as canonical isomeric SMILES.

Retained structures contained only B, Br, C, Cl, F, H, I, N, O, P, S, Se or Si; had more than eight heavy atoms; contained carbon as more than 20% of their heavy atoms; and had neutral-parent molecular weight below 800 Da. Isotopically labelled structures, unsupported valence or formal-charge states, and molecules failing RDKit parsing or sanitization were excluded.

All logarithms are base 10. Solubility was converted to mol/L before transformation to log *S* = log_10_(mol/L), whereas mass concentrations were divided by the standardized neutral-parent molecular weight before transformation. Permeability was converted to cm/s before transformation to log *P*_app_ = log_10_(cm/s). Efflux was represented as log_10_[*P*_app_(B *→* A)*/P*_app_(A *→* B)], and potency as p*X* = *−* log_10_(*X* in mol/L). Reported logP, and log *D*_7.4_ values were already logarithmic and dimensionless.

For duplicate global-ADMET records, measurements were first converted to the common endpoint unit. For exactly two measurements, their absolute difference was required to be below twice the assay-level standard deviation; otherwise, the group was excluded. Accepted pairs retained the measurement in the prespecified less-favorable direction—for example, lower solubility or permeability and higher efflux—to limit false-positive labels. For three or more measurements, groups were retained only when their within-group standard deviation was below 0.5 log units, and the observed value closest to the median was selected. Missing endpoints were never imputed.

Publication or temporal cohort membership was assigned before campaign analysis. Every submitted proposal remained in the molecular-validity denominator. Invalid proposals were excluded from fingerprints, descriptors, and model analyses but were recorded explicitly.

### S3.6 Information availability and leakage controls

The temporal split prevents explicit future endpoint values from being supplied to MAGI but cannot exclude exposure through public literature or model pretraining. Sanofi project-model training and future cohorts had no identical InChIKey overlap with a non-missing primary endpoint. For GSK, exact endpoint-labelled overlap with the global ADMET training collection was asserted to be zero.

To contextualize structure-conditioned predictions, one crystallographic reference complex was selected per target (Table S4). Successfully cofolded MAGI proposals were compared with the selected crystal ligand using residue-level interaction fingerprints.

**Table S4:**
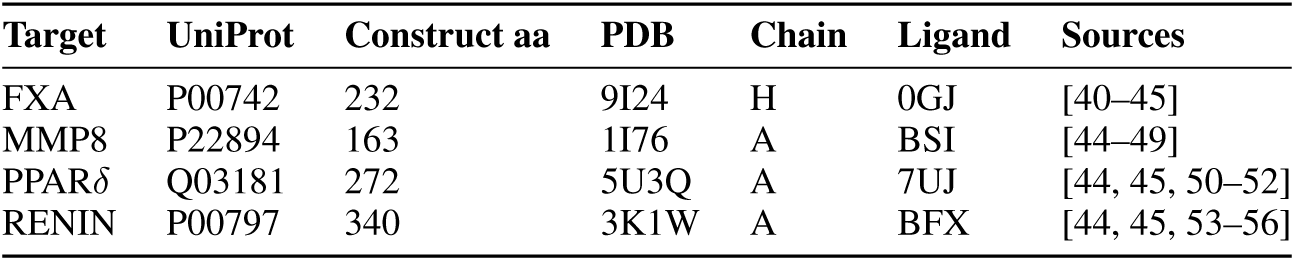
Protein constructs and selected crystallographic reference complexes used for structure-conditioned evaluation.

Interaction similarity was the Jaccard overlap between residue-level contact fingerprints for a generated pose and its target’s selected crystallographic ligand. Buffer components, ions, and incomplete ligands were excluded from the reference complexes. This analysis measures recovery of the selected binding interaction pattern [45]. MMP8 proposals showed the highest interaction-fingerprint similarity to the selected crystallographic ligand, followed by PPAR*δ* and RENIN, while FXA showed lower overlap (Figure S2b).

## S4 Services

### S4.1 Global ADMET models

The general ADMET ensemble was trained on 156,797 curated molecules carrying 186,120 sparse labels across seven endpoints. XGBoost models use RDKit physicochemical descriptors concatenated with 2,048-bit Avalon fingerprints [57]. Chemprop-MTL uses a depth-3 directed message-passing network with hidden dimension 300, a two-layer 300-unit FFN, dropout 0.2 and norm aggregation [58]. CheMeleon-MTL uses the foundation-initialized 2,048-dimensional, depth-6 message-passing network with a task-specific FFN and native aggregation [59, 60]. The multitask networks use MAE in standardized target space, inverse-standard-deviation task weighting, a 200-epoch cap, and early stopping patience 25. Pooled out-of-fold performance uses five shared Butina folds; deployed models were retrained on all data. The consensus is the mean across model families and cross-family standard deviation reports of disagreement. Performances are reported in Table S5.

**Figure S2:**
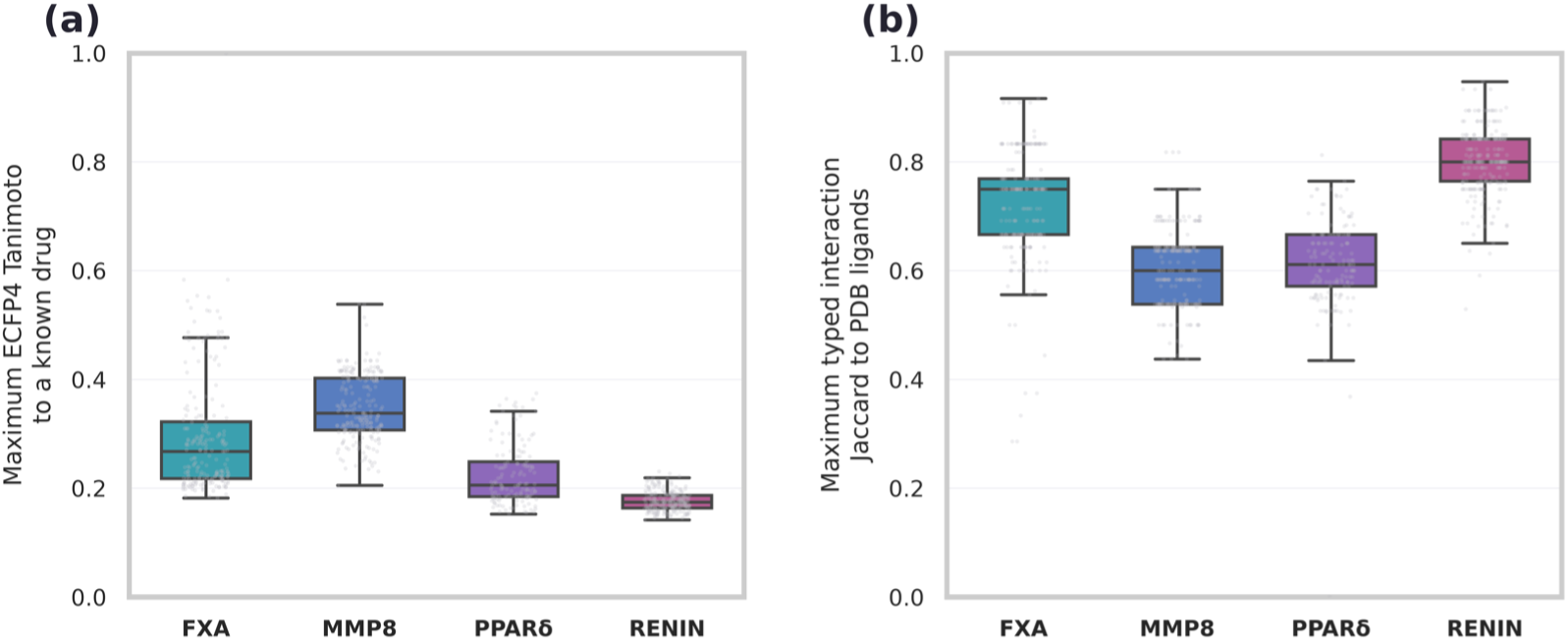
Similarity between the held-out set and publicly available data (a) ECFP4 similarity to known drugs. For each FXa, MMP8, PPAR*δ*, and RENIN held-out compound, points show the maximum ECFP4 Tanimoto similarity to a standardized drugs structures extracted from SureChEMBL. **(b) Interaction-fingerprint similarity to crystallographic ligands.** For each FXa, MMP8, PPAR*δ*, and RENIN held-out compound, points show the maximum interaction fingerprint with Jaccard similarity to available experimental structures extracted from the PDB. Boxes show medians and interquartile ranges, with whiskers extending to 1.5 times the interquartile range.

**Table S5:**
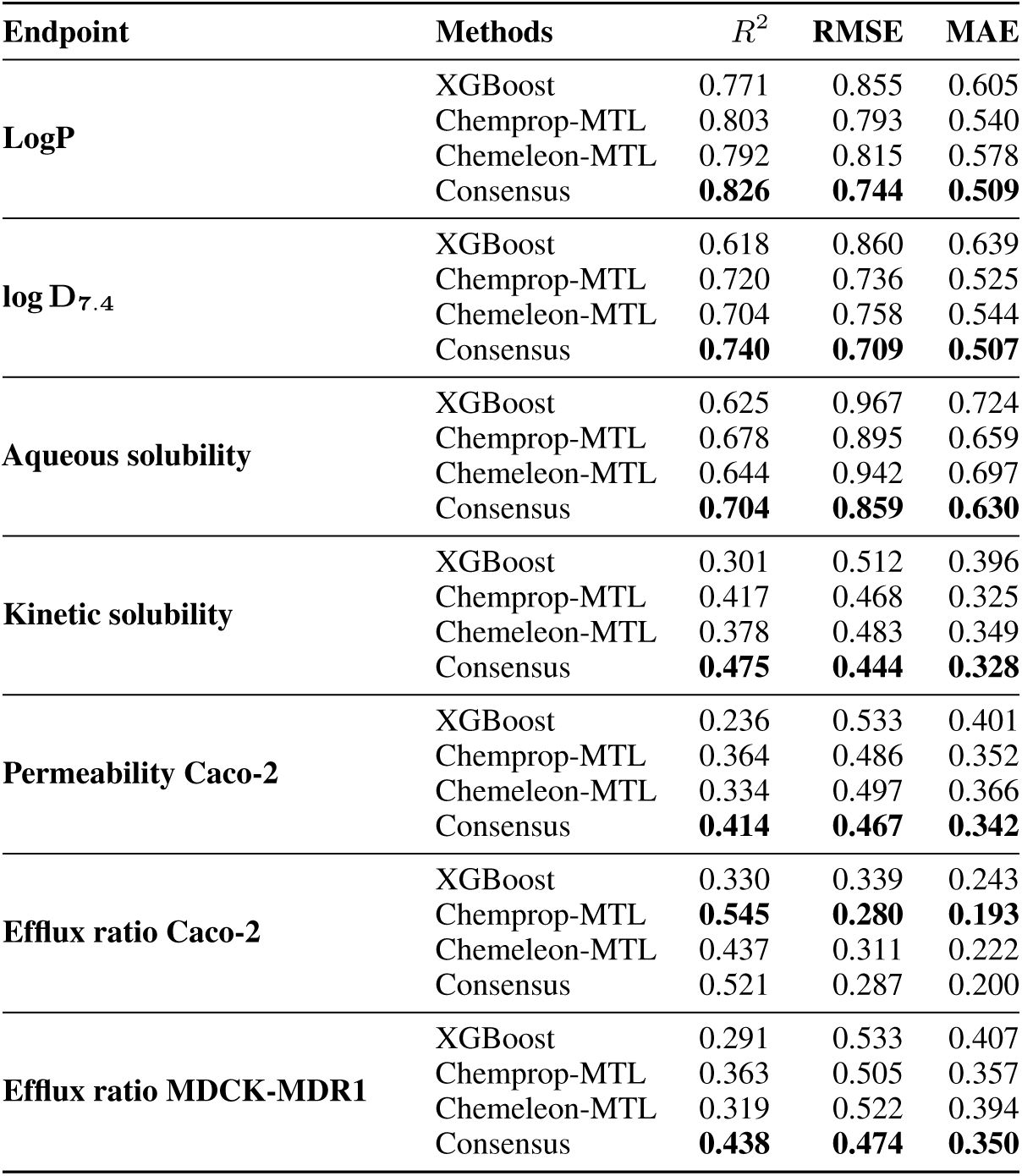
Pooled out-of-fold performance on five shared Butina folds. Metrics are calculated in each endpoint’s native transformed unit; *R*^2^ is unitless.

The trained ensemble was applied to a one-million-compound Enamine REAL sample to calibrate operational similarity thresholds. The fifth-percentile thresholds are Avalon nearest-neighbour Tanimoto *≥*0.4358 and Chemprop latent-space cosine *≥*0.7334. The deployed overall AD pass requires both criteria; per-axis scores and pass flags remain available. These thresholds indicate similarity to model training space and are not calibrated probabilities of prediction correctness. Held-out endpoint calibration is shown in Figure S3a.

The models are exposed through a persistent REST service that standardizes and scores each submitted SMILES and returns family predictions, consensus mean, cross-model disagreement and applicability metadata. The released inference artifact does not contain row-level endpoint labels, so exact endpoint-specific GSK training overlap is not claimed.

### S4.2 Local project-specific models

#### Project model recipes

Project-specific ligand-based models were developed only for the Sanofi campaigns as these campaigns included affinity endpoints supported by sufficiently large quantity of historical measurements for model training. Their primary purpose was to provide campaign-specific predictions for scoring and prioritization; they also provided a ligand-based baseline orthogonal to Boltz-2. Per endpoint we trained an ensemble of XGBoost, Chemprop MTL and CheMeleon MTL models; in the multitask objective the project endpoints carry half of the total task weight and the general ADMET endpoints enter as auxiliary tasks summing to the other half, up weighting the scarcer project data (fewer endpoints). Every model sees only compounds before the task temporal cutoff to prevent leakage and is served exactly like the global service. The target and temporal window must be fixed explicitly for a controlled replay; omitting the window causes the service to fall back to its latest snapshot. The ligand-based models were evaluated on the held-out sets shown in Figure S3b. The ligand-based project models reach rank correlations above 0.4 for FXA, MMP8 and PPAR*δ* (with low *R*^2^), while RENIN shows a low *R*^2^ at *ρ ≈* 0.31. Because no error-calibrated applicability-domain threshold was established for the project-specific models, cross-model disagreement is treated as a qualitative extrapolation warning and is not referred to as a formal out-of-domain classification.

### S4.3 Boltz-2

Boltz-2 predicts a protein–ligand complex and an affinity-derived pX for each submitted structure [32]. Sanofi task prompts designate this affinity as a co-primary objective and requires a PoseBusters-valid pose as a hard gate. Cofolding uses the fixed full target sequence and a precomputed MSA, with 3 recycling steps and 200 diffusion sampling steps (one diffusion sample per structure) followed by affinity sampling; the service returns the predicted complex, confidence score, ligand iPTM, affinity value and binder probability. The affinity output, log_10_(IC_50_*/*µM), is converted to the conventional p-scale. For throughput, the model is kept resident in GPU memory behind a persistent service, and Boltz-2 runs on a dedicated GPU while REINVENT 4 occupies a separate GPU.

The agreement with the ligand-based model on the held-out set was evaluated in Figure S3c. The two methods agree only for MMP8, whose project compounds have similar analogues in the PDB— despite different architectures and non-identical training sets, both encode similar information for that series. For FXA both models identify strong binders but disagree at the extremes (most- and least-active), giving poor overall agreement, while for PPAR*δ* and RENIN the predictions do not agree. This pattern reinforces that ligand- and structure-based predictions are complementary rather than redundant.

### S4.4 PoseBusters

PoseBusters screens the ligand extracted from each Boltz complex in mol mode, evaluating only ligand geometry and chemistry (bond lengths and angles, planarity, internal clashes, valences, etc.) without receptor-dependent docking validation. The score is the fraction of the 12 Boolean checks passed; all_pass requires every check and missing or invalid upstream structures to return null and are not imputed. Observed pass fractions range from 0.833 to 1.0, and a fraction below 1 is not reported as a complete pass. The same output is used in two ways: the pass fraction can enter the REINVENT objective as a structural-plausibility reward, and the per-check breakdown is returned to the agent as metadata flagging specific geometric/chemical liabilities.

**Figure S3:**
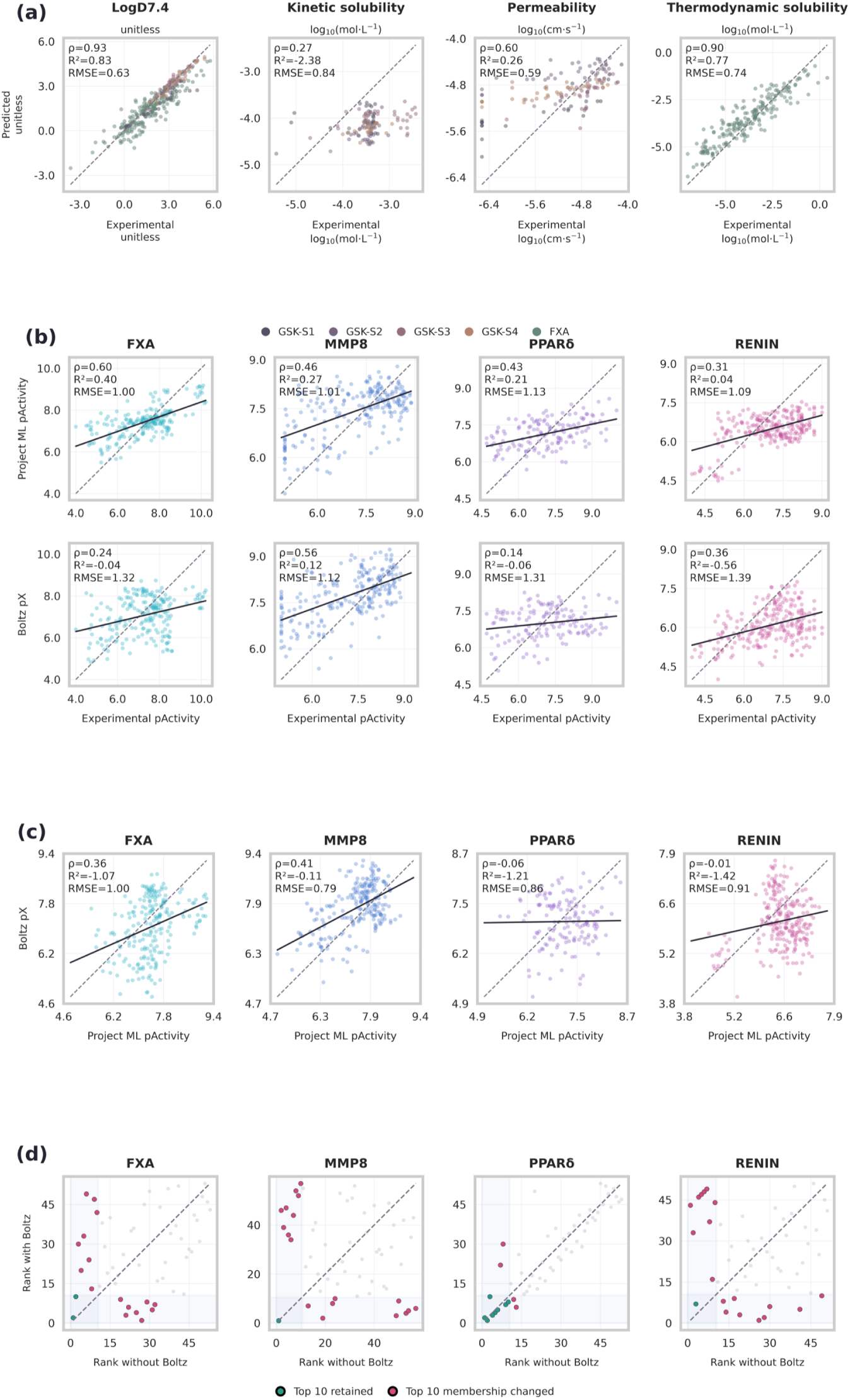
Validation of ligand-based models and Boltz-2 on experimental data. (a) Held-out validation of the ADMET models on the projects data. Scatter plots of the prediction from the consensus ADMET model to the GSK and FXA experimental ADMET data. Prediction are grouped by endpoints and colored by projects: GSK-S1 (light blue), GSK-S2 (purple), GSK-S3 (pink), GSK-S4 (orange), and FXA (green). (b) Predictive performance of Sanofi project models (top) and Boltz-2 affinity prediction (bottom) on the held-out set: experimental activity versus ligand-based model and Boltz-2 predictions for FXA, MMP8, PPAR*δ* and RENIN. Point are colored based on the target. (c) Agreement between ligand-based model and Boltz-2 affinity predictions on held-out Sanofi compounds: ligand-based model predictions versus Boltz-2 pX. (d) Post hoc sensitivity of proposal ranking to the Boltz-2 affinity term. Compounds were ranked with or without Boltz, evaluating ranking agreement between the two methods. Top 10 ranks with or without Boltz are covered by a grey area. Compounds with matching ranking are colored in green, and otherwise in red.

### S4.5 Interaction Profiler

The interaction profiler is an interpretation scoring service. In structure-plus-SMILES mode it (i) verifies ligand identity against the submitted SMILES, (ii) prepares and protonates both ligand and receptor at pH 7.4, (iii) crops an 8 Å binding-site region around the ligand, and (iv) enumerates protein–ligand contacts within 4 Å, returning typed contacts and a human readable interaction fingerprint, verifying medicinal chemistry taxonomy of molecule fragments. Each contact is recorded as a structured triplet linking: the ligand fragment named and localized under a medicinal-chemistry taxonomy; the interaction type (e.g., H-bond, hydrophobic, *π*-stacking, salt bridge, halogen bond) together with its distance, angle and geometric validity; the protein partner residue, the specific interacting atom, and whether it is a backbone or side chain. This mode can also be used in the objective function as an additional endpoint by reporting the count of distinct types of interactions. In SMILES-only mode it returns just the medicinal-chemistry fragment taxonomy.

## S5 Metrics

### S5.1 Attainment scope and limitations

Primary-endpoint attainment alone does not establish multiparameter success. We therefore report primary-objective attainment, secondary-objective attainment, fixed-guardrail compliance, and complete TTP attainment separately. A molecule attains the complete TTP only when all required objectives and guardrails pass.

For global ADMET models, predictions are classified as in domain, singly flagged, or outside the operational applicability domain according to the two fixed similarity rules defined in Section S4.1. Numerical predictions are retained for transparency but are flagged in analyses; attainment is summarized both overall and, where sample size permits, among in-domain predictions only. For Sanofi project-specific models, no formal applicability-domain classifier was implemented. Cross-model standard deviation is therefore reported as an extrapolation warning rather than used to classify a prediction as formally in or out of domain.

### S5.2 Molecular quality

Novelty is 1 minus the maximum Tanimoto similarity over ECFP4 fingerprints [37] to the shown set; internal diversity is 1 minus the mean pairwise Tanimoto; synthesizability is the synthetic-accessibility score [36]; scaffold provenance uses Bemis–Murcko frameworks [61]; all cheminformatics uses RDKit [62]. Calibration is predicted against measured agreement on held-out compounds, reported as Pearson *r*, mean absolute error and bias at the withheld optimized compound. The cofolding ablation re-ranks the same cofolded molecules by the ligand-only objective and by the objective including the Boltz-2 term, comparing orderings by Spearman rank correlation and top K overlap.

### S5.3 GSK and AstraZeneca primary-endpoint outcomes

Figure 1b reports ligand-model primary-endpoint distributions for the four GSK projects and the AstraZeneca campaign. Three of four GSK project produced at least one proposal satisfying the model-space primary threshold: REINVENT for GSK-S2 and both routes for GSK-S3 and GSK-S4. Neither route met the GSK-S1 kinetic-solubility objective, and LLM remained below the GSK-S2 permeability criterion. GSK-S1 held-out agreement was weak and range-compressed (Table S5; Figure S3a), making predictions near its target extrapolative. The agent consequently treated aqueous solubility as a secondary objective, where improvement was clearer (Figure S12).

### S5.4 Statistical analysis

For Figure 1a, each metric is averaged across rounds 1–5 within each project and route; LLM and REINVENT project means are compared with a two-sided Wilcoxon signed-rank test (four paired projects per organization), with Holm adjustment across the predefined tests. Project-round points remain visible as descriptive data. Within-project molecule-level Mann–Whitney comparisons in Figure 1b–c and Figures S4 and S9 are exploratory: canonical structures are deduplicated, but analogues remain chemically dependent, so their p-values are not interpreted as independent cross-project evidence. Chemist-evaluation proportions (Figure S18) are reported with Wilson 95% intervals.

## S6 Validation protocol

### S6.1 Project objectives

Each task provides a starter with measured ADMET, four measured reference analogues, an endpoint objective as a delta on the starter, and guardrails; potency is out of scope. S1: solubility from 330 µM (log *−*3.48), raise by at least 0.6 log and permeability to at least 100 nm/s, with CHI/log *D*_7.4_ below 3.5 and MW below 440 [27]. S2: apparent permeability from 17 nm/s (log *−*5.77), raise by at least 1.25 log, with solubility at least 150 µM, no added hydrogen-bond donors and MW below 470 [29]. S3: log *D*_7.4_ from 4.3, lower by at least 1.1 log, with solubility at least 1000 µM, permeability at least 150 nm/s and MW below 420 [28]. S4: log *D*_7.4_ from 5.7, lower by at least 2.85 log, with solubility at least 265 µM, permeability at least 100 nm/s and MW below 470 [30]. AZ: log *D*_7.4_ lowered below 3.0, with rotatable bonds at most 8. The agent receives the starter and analogue measurements, and never saw the held-out compounds.

**Table S6:** Project objectives, guardrails, scoring directions, target bounds, and required evaluations. Upper desirability values for increasing potency objectives are saturation points rather than exclusion thresholds.

| Project | Requirement | Endpoint or constraint | Direction | Minimum acceptable value | Desirability saturation value | Unit |
| --- | --- | --- | --- | --- | --- | --- |
| GSK-S1 | Primary objective | Kinetic solubility | Increase | −2.881 | − | $\log_{10}(\text{mol/L})$ |
| GSK-S1 | Fixed guardrail | Permeability | Minimum | −5.0 | − | $\log_{10}(\text{cm/s})$ |
| GSK-S1 | Fixed guardrail | $\log D_{7.4}$ | Maximum | − | 3.5 | $\log_{10}(D)$ |
| GSK-S1 | Fixed guardrail | Molecular weight | Maximum | − | 440.0 | g/mol |
| GSK-S2 | Primary objective | Permeability | Increase | −4.52 | − | $\log_{10}(\text{cm/s})$ |
| GSK-S2 | Fixed guardrail | Aqueous solubility | Minimum | 150.0 | − | $\mu\text{M}$ |
| GSK-S2 | Fixed guardrail | Hydrogen-bond donors | No increase | − | − | count |
| GSK-S2 | Fixed guardrail | Molecular weight | Maximum | − | 470.0 | g/mol |
| GSK-S3 | Primary objective | $\log D_{7.4}$ | Decrease | − | 3.2 | $\log_{10}(D)$ |
| GSK-S3 | Fixed guardrail | Aqueous solubility | Minimum | 1000.0 | − | $\mu\text{M}$ |
| GSK-S3 | Fixed guardrail | Permeability | Minimum | 150.0 | − | nm/s |
| GSK-S3 | Fixed guardrail | Molecular weight | Maximum | − | 420.0 | g/mol |
| GSK-S4 | Primary objective | $\log D_{7.4}$ | Decrease | − | 2.85 | $\log_{10}(D)$ |
| GSK-S4 | Fixed guardrail | Aqueous solubility | Minimum | 265.0 | − | $\mu\text{M}$ |
| GSK-S4 | Fixed guardrail | Permeability | Minimum | 100.0 | − | nm/s |
| GSK-S4 | Fixed guardrail | Molecular weight | Maximum | − | 470.0 | g/mol |
| AZ | Primary objective | $\log D_{7.4}$ | Decrease | − | 3.0 | $\log_{10}(D)$ |
| AZ | Fixed guardrail | Rotatable bonds | Maximum | − | 8.0 | count |
| AZ | Fixed guardrail | Molecular weight | In range | 250.0 | 475.0 | g/mol |
| FXA | Primary objective | $\text{p}K_i$ | Increase | 8.5 | 9.0 | p-scale |
| FXA | Secondary objective | Aqueous solubility | Increase | −4.301 | −4.0 | $\log_{10}(\text{mol/L})$ |
| FXA | Secondary objective | $\log D_{7.4}$ | In range | 1.5 | 3.5 | $\log_{10}(D)$ |
| FXA | Fixed guardrail | Molecular weight | In range | 400.0 | 675.0 | g/mol |
| FXA | Required evaluation | Boltz-2 affinity and PoseBusters validity | Required | − | − | p-scale |
| MMP8 | Primary objective | $\text{pIC}_{50}$ | Increase | 8.5 | 9.0 | p-scale |
| MMP8 | Fixed guardrail | Molecular weight | In range | 350.0 | 575.0 | g/mol |
| MMP8 | Required evaluation | Boltz-2 affinity and PoseBusters validity | Required | − | − | p-scale |
| PPARδ | Primary objective | $\text{pEC}_{50}$ | Increase | 8.0 | 9.0 | p-scale |
| PPARδ | Fixed guardrail | Molecular weight | In range | 425.0 | 625.0 | g/mol |
| PPARδ | Required evaluation | Boltz-2 affinity and PoseBusters validity | Required | – | – | p-scale |
| RENIN | Primary objective | pIC<sub>50</sub> | Increase | 8.0 | 9.0 | p-scale |
| RENIN | Fixed guardrail | Molecular weight | In range | 325.0 | 525.0 | g/mol |
| RENIN | Required evaluation | Boltz-2 affinity and PoseBusters validity | Required | – | – | p-scale |

### S6.2 Agent experiment

The LLM route asks Claude Sonnet 4.6 to return approximately five final structures per round. The REINVENT route creates or updates a TTP, invokes REINVENT 4, monitors or recovers the run, retrieves its pool, and selects five final structures. All reported route figures and statistics use the common complete rounds 1–5 window. GSK and AstraZeneca experiments used the same ADMET evaluators and exclude cofolding. Sanofi prompts additionally designate project-model prediction and Boltz-2 affinity as co-primary, a PoseBusters-valid pose as a hard gate, and interaction analysis as informative metadata.

**Table S7:** Project-level REINVENT generation pool, shortlist and cofold coverage.

| Project | Pool records n | Valid distinct structures | Unique shortlisted | Shortlist construction | Cofolded structures | Cofold fraction |
| --- | --- | --- | --- | --- | --- | --- |
| GSK-S1 | 96061 | 94363 | 25 | Agent-authored top five list | – | – |
| GSK-S2 | 132765 | 131030 | 25 | Agent-authored top five list | – | – |
| GSK-S3 | 200977 | 195351 | 20 | Agent-authored top five list | – | – |
| GSK-S4 | 204769 | 198243 | 25 | Agent-authored top five list | – | – |
| AZ | 163840 | – | 25 | Agent-authored top five list | – | – |
| FXA | 63455 | 63455 | 21 | Agent-authored final top five from transcript | 61260 | 0.965 |
| MMP8 | 117061 | 117061 | 22 | Agent-authored final top five from transcript | 114250 | 0.976 |
| PPARδ | 29328 | 29328 | 18 | Agent-authored final top five from transcript | 28345 | 0.966 |
| RENIN | 74489 | 74489 | 25 | Agent-authored final top five from transcript | 68266 | 0.916 |

**Table S8:** Exact per-invocation of REINVENT parameters parsed from audited operation records.

| Project | Round | Invocation ID<br>in round | Maximum steps | Batch size | Sigma | Seed<br>structures<br>n |
| --- | --- | --- | --- | --- | --- | --- |
| GSK-S1 | 1 | 1 | 75 | 64 | 120 | 5 |
| GSK-S1 | 2 | 1 | 80 | 64 | 130 | 5 |
| GSK-S1 | 3 | 1 | 80 | 64 | 130 | 5 |
| GSK-S1 | 4 | 1 | 80 | 64 | 120 | 5 |
| GSK-S1 | 5 | 1 | 80 | 64 | 115 | 6 |
| GSK-S3 | 1 | 1 | 75 | 96 | 120 | 5 |
| GSK-S3 | 2 | 1 | 85 | 96 | 140 | 5 |
| GSK-S3 | 3 | 1 | 90 | 96 | 130 | 5 |
| GSK-S3 | 4 | 1 | 80 | 96 | 140 | 5 |
| GSK-S3 | 5 | 1 | 75 | 96 | 140 | 5 |
| GSK-S2 | 1 | 1 | 75 | 64 | 120 | 5 |
| GSK-S2 | 2 | 1 | 80 | 64 | 128 | 5 |
| GSK-S2 | 3 | 1 | 90 | 64 | 140 | 5 |
| GSK-S2 | 4 | 1 | 90 | 64 | 150 | 5 |
| GSK-S2 | 5 | 1 | 90 | 64 | 160 | 5 |
| GSK-S4 | 1 | 1/2 | 75/75 | 64/64 | 120/120 | 5 |
| GSK-S4 | 2 | 1 | 80 | 64 | 130 | 5 |
| GSK-S4 | 3 | 1/2 | 80/80 | 64/64 | 130/130 | 5 |
| GSK-S4 | 4 | 1 | 80 | 64 | 130 | 5 |
| GSK-S4 | 5 | 1 | 80 | 64 | 130 | 5 |
| AZ | 1 | 1 | 80 | 384 | 128 | 5 |
| AZ | 2 | 1 | 80 | 704 | 128 | 5 |
| AZ | 3 | 1 | 100 | 384 | 128 | 5 |
| AZ | 4 | 1 | 100 | 256 | 128 | 5 |
| AZ | 5 | 1 | 100 | 128 | 128 | 5 |
| FXA | 1 | 1/2 | 300/300 | 64/64 | 120/128 | 5 |
| FXA | 2 | 1 | 300 | 320 | 120 | 2 |
| FXA | 3 | 1/2 | 300/300 | 64/64 | 128/128 | 2 |
| FXA | 4 | 1 | 300 | 64 | 120 | 5 |
| FXA | 5 | 1 | 300 | 64 | 120 | 5 |
| MMP8 | 1 | 1/2/3 | 200/200/200 | 32/32/32 | 120/120/120 | 5 |
| MMP8 | 2 | 1/2 | 300/300 | 128/128 | 128/128 | 5 |
| MMP8 | 3 | 1 | 250 | 64 | 120 | 5 |
| MMP8 | 4 | 1 | 150 | 64 | 120 | 6 |
| MMP8 | 5 | 1 | 200 | 128 | 128 | 5 |
| PPARδ | 1 | 1 | 200 | 64 | 120 | 5 |
| PPARδ | 2 | 1/2 | 250/250 | 64/32 | 120/120 | 5 |
| PPARδ | 2 | 2 | 250 | 32 | 120 | 5 |
| PPARδ | 3 | 1 | 300 | 64 | 120 | 5 |
| PPARδ | 4 | 1/2/3 | 300/300/300 | 64/64/32 | 120/120/120 | 6 |
| PPARδ | 5 | 1 | 300 | 80 | 120 | 2 |
| RENIN | 1 | 1/2/3 | 200/200/200 | 32/32/32 | 120/120/120 | 5 |
| RENIN | 2 | 1/2/3 | 200/200/200 | 128/128/128 | 120/120/120 | 5 |
| RENIN | 3 | 1 | 200 | 64 | 120 | 3 |
| RENIN | 4 | 1/2 | 250/250 | 128/128 | 110/110 | 5 |
| RENIN | 5 | 1/2 | 300/300 | 96/96 | 120/120 | 5 |

### S6.3 Medicinal chemist evaluation

The prepared blinded instrument contains 28 questions: 12 source-discrimination cases, 11 structure-first candidate-prioritization cases, and 5 SAR-reasoning cases. Source-discrimination cases ask which of one agent’s proposal, and one similarity-matched held-out compound is AI-generated. Prioritization cases ask the evaluator to select among blinded LLM, REINVENT, and held-out candidates under an explicit property objective. SAR cases request a verdict of supported, plausible but incomplete, or unsupported, together with confidence and an optional negative-reason code.

Primary analyses are discrimination accuracy with a binomial 95% interval relative to chance, candidate-choice proportions, and reasoning-verdict proportions. Results for the four chemists who completed the instrument are shown in Figure S18. Discrimination accuracy did not separate from chance for any chemist or in pooled analysis, and no source was preferentially selected in the prioritization cases.

## S7 Availability and ethics

### S7.1 Human-participant ethics

The blinded medicinal-chemist evaluation involved 4 participants. No personally identifying participant information is released.

### S7.2 Responsible release

Molecular design systems have potential dual-use applications and may generate unsafe, reactive, or otherwise inappropriate structures. MAGI outputs are predictions rather than experimentally validated compounds and require expert review.

## S8 Supplementary figures

Extended figures supporting the main text analyses are shown below. Main-text cross-references identify the relevant panels; underlying numerical values and tests are provided in the accompanying data package.

## NeurIPS Paper Checklist

### 1. Claims

Question: Do the main claims made in the abstract and introduction accurately reflect the paper’s contributions and scope?

Answer: [Yes]

Justification: The three contributions listed in Section 1 map directly onto the reported results: the versioned executable objective shared by both generation routes (Section 3.2), the campaign-level evaluation on nine industry-derived tasks (Sections 5.1–5.3), and the analysis of evidence-driven adaptation (Section 5.3 and Section S2.6). The abstract states that attainment was governed primarily by predictor calibration rather than generation route, which is the conclusion supported by Section 5.2.

Guidelines:

- The answer [N/A] means that the abstract and introduction do not include the claims made in the paper.
- The abstract and/or introduction should clearly state the claims made, including the contributions made in the paper and important assumptions and limitations. A [No] or [N/A] answer to this question will not be perceived well by the reviewers.
- The claims made should match theoretical and experimental results, and reflect how much the results can be expected to generalize to other settings.
- It is fine to include aspirational goals as motivation as long as it is clear that these goals are not attained by the paper.

### 2. Limitations

Question: Does the paper discuss the limitations of the work performed by the authors?

Answer: [Yes]

Justification: Section 7 is a dedicated Limitations section. It states that all proposal endpoints are predictions, so historical measurements represent a different evidence level; that each condition used one reasoning backend and one trajectory; and that temporal splits cannot exclude pretraining exposure.

Guidelines:

- The answer [N/A] means that the paper has no limitation while the answer [No] means that the paper has limitations, but those are not discussed in the paper.
- The authors are encouraged to create a separate “Limitations” section in their paper.
- The paper should point out any strong assumptions and how robust the results are to violations of these assumptions (e.g., independence assumptions, noiseless settings, model well-specification, asymptotic approximations only holding locally). The authors should reflect on how these assumptions might be violated in practice and what the implications would be.
- The authors should reflect on the scope of the claims made, e.g., if the approach was only tested on a few datasets or with a few runs. In general, empirical results often depend on implicit assumptions, which should be articulated.
- The authors should reflect on the factors that influence the performance of the approach. For example, a facial recognition algorithm may perform poorly when image resolution is low or images are taken in low lighting. Or a speech-to-text system might not be used reliably to provide closed captions for online lectures because it fails to handle technical jargon.
- The authors should discuss the computational efficiency of the proposed algorithms and how they scale with dataset size.
- If applicable, the authors should discuss possible limitations of their approach to address problems of privacy and fairness.
- While the authors might fear that complete honesty about limitations might be used by reviewers as grounds for rejection, a worse outcome might be that reviewers discover limitations that aren’t acknowledged in the paper. The authors should use their best judgment and recognize that individual actions in favor of transparency play an important role in developing norms that preserve the integrity of the community. Reviewers will be specifically instructed to not penalize honesty concerning limitations.

### 3. Theory assumptions and proofs

Question: For each theoretical result, does the paper provide the full set of assumptions and a complete (and correct) proof?

Answer: [N/A]

Justification: The paper contains no theoretical results. Equations (S1) and (S2) are definitions of the endpoint desirability transform and the weighted Derringer–Suich aggregation adopted from prior work [35], not claims requiring proof.

Guidelines:

- The answer [N/A] means that the paper does not include theoretical results.
- All the theorems, formulas, and proofs in the paper should be numbered and cross-referenced.
- All assumptions should be clearly stated or referenced in the statement of any theorems.
- The proofs can either appear in the main paper or the supplemental material, but if they appear in the supplemental material, the authors are encouraged to provide a short proof sketch to provide intuition.
- Inversely, any informal proof provided in the core of the paper should be complemented by formal proofs provided in appendix or supplemental material.
- Theorems and Lemmas that the proof relies upon should be properly referenced.

### 4. Experimental result reproducibility

Question: Does the paper fully disclose all the information needed to reproduce the main experimental results of the paper to the extent that it affects the main claims and/or conclusions of the paper (regardless of whether the code and data are provided or not)?

Answer: [Yes]

Justification: Section S2 specifies the agent architecture, the TTP schema and the tool loop; Section S3 specifies dataset curation, standardization and the temporal cutoffs; Section S4 specifies every model architecture and hyperparameter; and Section S6 specifies the campaign protocol. Table S1 lists deployed service versions, Table S6 the per-project objectives and guardrails, and Table S8 the exact per-invocation REINVENT parameters. The reasoning backend is identified in Section S2.5, which also notes that provider-side sampling defaults were used and no seed was fixed.

Guidelines:

- The answer [N/A] means that the paper does not include experiments.
- If the paper includes experiments, a [No] answer to this question will not be perceived well by the reviewers: Making the paper reproducible is important, regardless of whether the code and data are provided or not.
- If the contribution is a dataset and/or model, the authors should describe the steps taken to make their results reproducible or verifiable.
- Depending on the contribution, reproducibility can be accomplished in various ways. For example, if the contribution is a novel architecture, describing the architecture fully might suffice, or if the contribution is a specific model and empirical evaluation, it may be necessary to either make it possible for others to replicate the model with the same dataset, or provide access to the model. In general. releasing code and data is often one good way to accomplish this, but reproducibility can also be provided via detailed instructions for how to replicate the results, access to a hosted model (e.g., in the case of a large language model), releasing of a model checkpoint, or other means that are appropriate to the research performed.
- While NeurIPS does not require releasing code, the conference does require all submissions to provide some reasonable avenue for reproducibility, which may depend on the nature of the contribution. For example
- If the contribution is primarily a new algorithm, the paper should make it clear how to reproduce that algorithm.
- If the contribution is primarily a new model architecture, the paper should describe the architecture clearly and fully.
- If the contribution is a new model (e.g., a large language model), then there should either be a way to access this model for reproducing the results or a way to reproduce the model (e.g., with an open-source dataset or instructions for how to construct the dataset).
- We recognize that reproducibility may be tricky in some cases, in which case authors are welcome to describe the particular way they provide for reproducibility. In the case of closed-source models, it may be that access to the model is limited in some way (e.g., to registered users), but it should be possible for other researchers to have some path to reproducing or verifying the results.

### 5. Open access to data and code

Question: Does the paper provide open access to the data and code, with sufficient instructions to faithfully reproduce the main experimental results, as described in supplemental material?

Answer: [Yes]

Justification: The MAGI agent, its MCP server and the global ADMET ensemble will be released under an open-source licence upon publication, with instructions covering environment setup, service contracts and the commands needed to replay a campaign. The RetroDMTA projects are third-party assets released under [31] with an accompanying code DOI, and the GSK series are extracted from published articles [27–30]. The AstraZeneca campaign data are not releasable; results depending on them are therefore not reproducible from released artefacts alone.

Guidelines:

- The answer [N/A] means that paper does not include experiments requiring code.
- Please see the NeurIPS code and data submission guidelines (https://neurips.cc/ public/guides/CodeSubmissionPolicy) for more details.
- While we encourage the release of code and data, we understand that this might not be possible, so [No] is an acceptable answer. Papers cannot be rejected simply for not including code, unless this is central to the contribution (e.g., for a new open-source benchmark).
- The instructions should contain the exact command and environment needed to run to reproduce the results. See the NeurIPS code and data submission guidelines (https://neurips.cc/public/guides/CodeSubmissionPolicy) for more details.
- The authors should provide instructions on data access and preparation, including how to access the raw data, preprocessed data, intermediate data, and generated data, etc.
- The authors should provide scripts to reproduce all experimental results for the new proposed method and baselines. If only a subset of experiments are reproducible, they should state which ones are omitted from the script and why.
- At submission time, to preserve anonymity, the authors should release anonymized versions (if applicable).
- Providing as much information as possible in supplemental material (appended to the paper) is recommended, but including URLs to data and code is permitted.

### 6. Experimental setting/details

Question: Does the paper specify all the training and test details (e.g., data splits, hyperparameters, how they were chosen, type of optimizer) necessary to understand the results?

Answer: [Yes]

Justification: Section S4.1 gives the ADMET model architectures, loss, task weighting, epoch cap and early-stopping patience, and the five shared Butina folds; Section S4.2 gives the project-model recipe and temporal restriction; Section S4.3 gives the recycling and diffusion sampling settings. Splits and cohort sizes are in Table S3 and Section S3.4, and per-invocation generation hyperparameters in Table S8.

Guidelines:

- The answer [N/A] means that the paper does not include experiments.
- The experimental setting should be presented in the core of the paper to a level of detail that is necessary to appreciate the results and make sense of them.
- The full details can be provided either with the code, in appendix, or as supplemental material.

### 7. Experiment statistical significance

Question: Does the paper report error bars suitably and correctly defined or other appropriate information about the statistical significance of the experiments?

Answer: [Yes]

Justification: Section S5.4 defines the experimental unit (project) and the repeated measure (round), and specifies two-sided Wilcoxon signed-rank tests over four paired projects per organization with Holm adjustment for Figure 1a. Box plots report medians, interquartile ranges and 1.5-IQR whiskers, and leakage metrics use bootstrap 95% intervals (Section S5.2). Section S5.4 explicitly labels the within-project molecule-level Mann–Whitney comparisons as exploratory because analogues remain chemically dependent.

Guidelines:

- The answer [N/A] means that the paper does not include experiments.
- The authors should answer [Yes] if the results are accompanied by error bars, confidence intervals, or statistical significance tests, at least for the experiments that support the main claims of the paper.
- The factors of variability that the error bars are capturing should be clearly stated (for example, train/test split, initialization, random drawing of some parameter, or overall run with given experimental conditions).
- The method for calculating the error bars should be explained (closed form formula, call to a library function, bootstrap, etc.)
- The assumptions made should be given (e.g., Normally distributed errors).
- It should be clear whether the error bar is the standard deviation or the standard error of the mean.
- It is OK to report 1-sigma error bars, but one should state it. The authors should preferably report a 2-sigma error bar than state that they have a 96% CI, if the hypothesis of Normality of errors is not verified.
- For asymmetric distributions, the authors should be careful not to show in tables or figures symmetric error bars that would yield results that are out of range (e.g., negative error rates).
- If error bars are reported in tables or plots, the authors should explain in the text how they were calculated and reference the corresponding figures or tables in the text.

### 8. Experiments compute resources

Question: For each experiment, does the paper provide sufficient information on the computer resources (type of compute workers, memory, time of execution) needed to reproduce the experiments?

Answer: [Yes]

Justification: Section S2.7 reports the hardware (AMD EPYC 9575F CPUs, AMD Instinct MI355X accelerators, ROCm 7.0), states that REINVENT and Boltz-2 occupied separate GPUs and that the remaining services were CPU-parallelized. Generation workloads are in Tables S7 and S8, and Figure S17 reports input-token volume against observable operations.

Guidelines:

- The answer [N/A] means that the paper does not include experiments.
- The paper should indicate the type of compute workers CPU or GPU, internal cluster, or cloud provider, including relevant memory and storage.
- The paper should provide the amount of compute required for each of the individual experimental runs as well as estimate the total compute.
- The paper should disclose whether the full research project required more compute than the experiments reported in the paper (e.g., preliminary or failed experiments that didn’t make it into the paper).

### 9. Code of ethics

Question: Does the research conducted in the paper conform, in every respect, with the NeurIPS Code of Ethics https://neurips.cc/public/EthicsGuidelines?

Answer: [Yes]

Justification: The research uses published medicinal-chemistry literature and de-identified industrial project data, releases no personally identifying information about the blinded evaluators (Section S7.1), and discusses the dual-use profile of molecular design in Section S7.2. Anonymity is preserved throughout the submission.

Guidelines:

- The answer [N/A] means that the authors have not reviewed the NeurIPS Code of Ethics.
- If the authors answer [No], they should explain the special circumstances that require a deviation from the Code of Ethics.
- The authors should make sure to preserve anonymity (e.g., if there is a special consideration due to laws or regulations in their jurisdiction).

### 10. Broader impacts

Question: Does the paper discuss both potential positive societal impacts and negative societal impacts of the work performed?

Answer: [Yes]

Justification: Section S7.2 states that molecular design systems have potential dual-use applications and may generate unsafe, reactive or otherwise inappropriate structures. The positive framing, accelerating lead optimization as an adaptive coordination layer over established models rather than replacing them, is developed in Sections 6 and 8.

Guidelines:

- The answer [N/A] means that there is no societal impact of the work performed.
- If the authors answer [N/A] or [No], they should explain why their work has no societal impact or why the paper does not address societal impact.
- Examples of negative societal impacts include potential malicious or unintended uses (e.g., disinformation, generating fake profiles, surveillance), fairness considerations (e.g., deployment of technologies that could make decisions that unfairly impact specific groups), privacy considerations, and security considerations.
- The conference expects that many papers will be foundational research and not tied to particular applications, let alone deployments. However, if there is a direct path to any negative applications, the authors should point it out. For example, it is legitimate to point out that an improvement in the quality of generative models could be used to generate Deepfakes for disinformation. On the other hand, it is not needed to point out that a generic algorithm for optimizing neural networks could enable people to train models that generate Deepfakes faster.
- The authors should consider possible harms that could arise when the technology is being used as intended and functioning correctly, harms that could arise when the technology is being used as intended but gives incorrect results, and harms following from (intentional or unintentional) misuse of the technology.
- If there are negative societal impacts, the authors could also discuss possible mitigation strategies (e.g., gated release of models, providing defenses in addition to attacks, mechanisms for monitoring misuse, mechanisms to monitor how a system learns from feedback over time, improving the efficiency and accessibility of ML).

### 11. Safeguards

Question: Does the paper describe safeguards that have been put in place for responsible release of data or models that have a high risk for misuse (e.g., pre-trained language models, image generators, or scraped datasets)?

Answer: [Yes]

Justification: Section S7.2 states that MAGI outputs are predictions rather than experimentally validated compounds and require expert review. The system is an orchestration layer over existing published generators and predictors and introduces no new generative capability beyond REINVENT 4 and the configured reasoning backend, and the industrial project data are de-identified.

Guidelines:

- The answer [N/A] means that the paper poses no such risks.
- Released models that have a high risk for misuse or dual-use should be released with necessary safeguards to allow for controlled use of the model, for example by requiring that users adhere to usage guidelines or restrictions to access the model or implementing safety filters.
- Datasets that have been scraped from the Internet could pose safety risks. The authors should describe how they avoided releasing unsafe images.
- We recognize that providing effective safeguards is challenging, and many papers do not require this, but we encourage authors to take this into account and make a best faith effort.

### 12. Licenses for existing assets

Question: Are the creators or original owners of assets (e.g., code, data, models), used in the paper, properly credited and are the license and terms of use explicitly mentioned and properly respected?

Answer: [Yes]

Justification: Every external asset is cited with the deployed version in Table S1: REINVENT 4 [2], Boltz-2 [32], PoseBusters [33], RDKit [62], XGBoost [57], Chemprop [58], CheMeleon [60] and UMAP [63]. Data sources are credited as OCHEM [38], ChEMBL [39], the PDB [45], UniProt [44], the RetroDMTA benchmark [31] and the four published GSK series.

Guidelines:

- The answer [N/A] means that the paper does not use existing assets.
- The authors should cite the original paper that produced the code package or dataset.
- The authors should state which version of the asset is used and, if possible, include a URL.
- The name of the license (e.g., CC-BY 4.0) should be included for each asset.
- For scraped data from a particular source (e.g., website), the copyright and terms of service of that source should be provided.
- If assets are released, the license, copyright information, and terms of use in the package should be provided. For popular datasets, paperswithcode.com/datasets has curated licenses for some datasets. Their licensing guide can help determine the license of a dataset.
- For existing datasets that are re-packaged, both the original license and the license of the derived asset (if it has changed) should be provided.
- If this information is not available online, the authors are encouraged to reach out to the asset’s creators.

### 13. New assets

Question: Are new assets introduced in the paper well documented and is the documentation provided alongside the assets?

Answer: [Yes]

Justification: The paper introduces the MAGI agent, its MCP server, the external REST scoring component for REINVENT 4, the global ADMET ensemble and the interaction profiler. These are described in Sections S2 and S4. Code and the trained global ADMET ensemble will be released under an open-source licence upon publication, with documentation covering installation, service contracts, applicability-domain caveats and known limitations.

Guidelines:

- The answer [N/A] means that the paper does not release new assets.
- Researchers should communicate the details of the dataset/code/model as part of their submissions via structured templates. This includes details about training, license, limitations, etc.
- The paper should discuss whether and how consent was obtained from people whose asset is used.
- At submission time, remember to anonymize your assets (if applicable). You can either create an anonymized URL or include an anonymized zip file.

### 14. Crowdsourcing and research with human subjects

Question: For crowdsourcing experiments and research with human subjects, does the paper include the full text of instructions given to participants and screenshots, if applicable, as well as details about compensation (if any)?

Answer: [Yes]

Justification: The blinded chemist evaluation involves human subjects. Section S6.3 describes the instrument in full: 28 questions across three case types, the exact response format required for each, and the primary analyses. Participants were four practising medicinal chemists or computational chemists; the exercise formed part of their normal work and no separate compensation was offered. Section S7.1 states that no personally identifying participant information is released.

Guidelines:

- The answer [N/A] means that the paper does not involve crowdsourcing nor research with human subjects.
- Including this information in the supplemental material is fine, but if the main contribution of the paper involves human subjects, then as much detail as possible should be included in the main paper.
- According to the NeurIPS Code of Ethics, workers involved in data collection, curation, or other labor should be paid at least the minimum wage in the country of the data collector.

### 15. Institutional review board (IRB) approvals or equivalent for research with human subjects

Question: Does the paper describe potential risks incurred by study participants, whether such risks were disclosed to the subjects, and whether Institutional Review Board (IRB) approvals (or an equivalent approval/review based on the requirements of your country or institution) were obtained?

Answer: [No]

Justification: Section S7.1 states only that no personally identifying participant information is released. The study consists of domain experts rating blinded molecular proposals and carries no risk.

Guidelines:

- The answer [N/A] means that the paper does not involve crowdsourcing nor research with human subjects.
- Depending on the country in which research is conducted, IRB approval (or equivalent) may be required for any human subjects research. If you obtained IRB approval, you should clearly state this in the paper.
- We recognize that the procedures for this may vary significantly between institutions and locations, and we expect authors to adhere to the NeurIPS Code of Ethics and the guidelines for their institution.
- For initial submissions, do not include any information that would break anonymity (if applicable), such as the institution conducting the review.

### 16. Declaration of LLM usage

Question: Does the paper describe the usage of LLMs if it is an important, original, or non-standard component of the core methods in this research? Note that if the LLM is used only for writing, editing, or formatting purposes and does *not* impact the core methodology, scientific rigor, or originality of the research, declaration is not required.

Answer: [Yes]

Justification: An LLM is the core of the contribution, not a writing aid. Section 3.1 identifies Claude Sonnet 4.6 as the configured reasoning backend that authors and revises objectives, invokes tools and selects candidates, and one of the two evaluated generation routes has the LLM propose molecules directly. Section S2.5 describes the system prompt, states that provider-side sampling defaults were used with no fixed seed, and restricts the analysis to observable messages, tool requests, tool results and configuration payloads.

Guidelines:

- The answer [N/A] means that the core method development in this research does not involve LLMs as any important, original, or non-standard components.
- Please refer to our LLM policy in the NeurIPS handbook for what should or should not be described.

**Figure S4:**
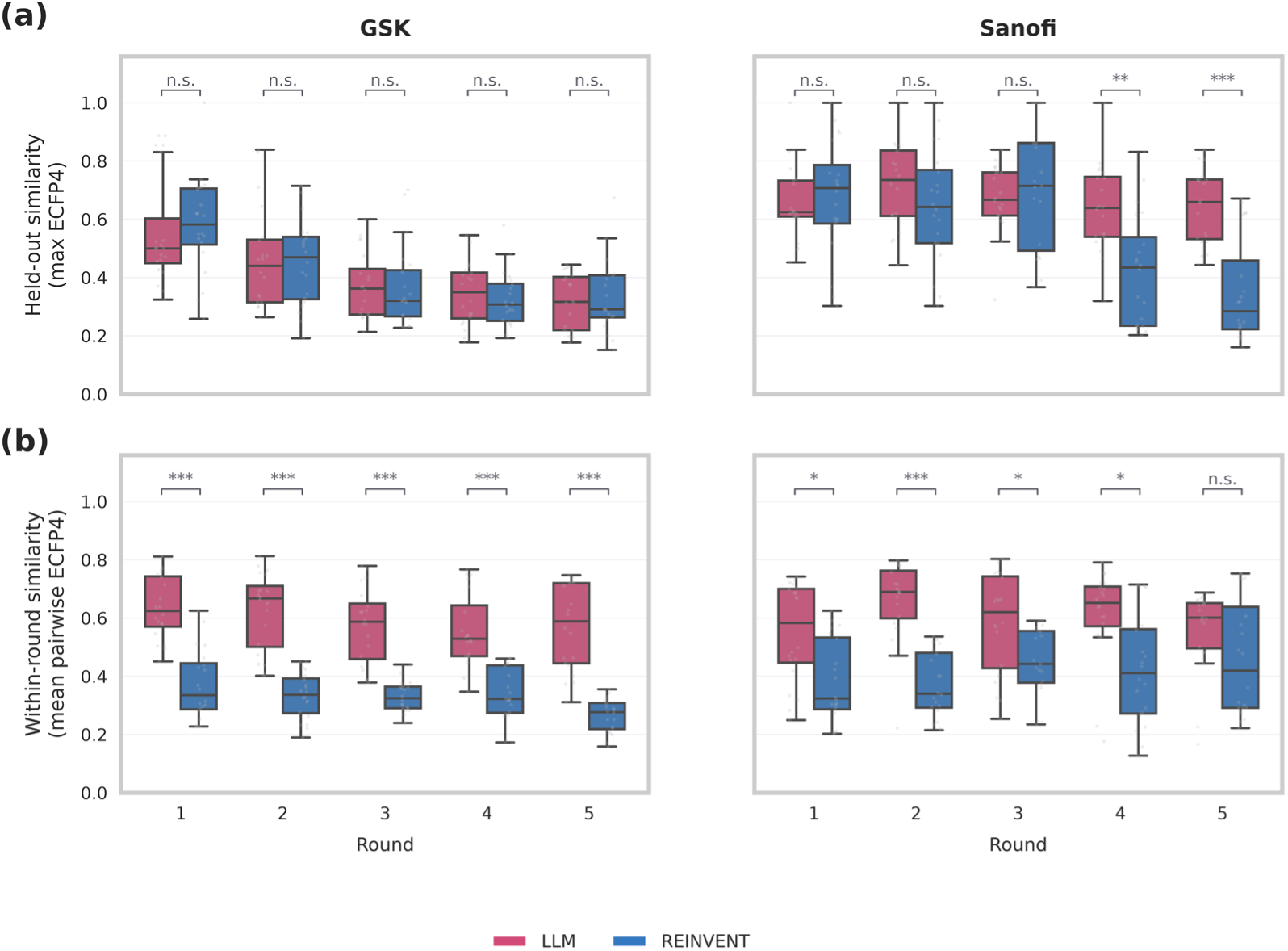
Round-by-round similarity to held-out chemistry and intra-round similarity. Red boxes represent LLM proposals, and blue boxes represent final agent-selected REINVENT proposals during rounds 1–5. Columns separate GSK and Sanofi projects. For each molecule, the upper row reports its maximum ECFP4 Tanimoto similarity to any project-specific held-out compound. The lower row reports its mean pairwise ECFP4 Tanimoto similarity to other molecules from the same project, route, and round; lower values therefore indicate greater intra-round diversity. Boxes show medians and interquartile ranges; individual outliers are not shown. Brackets compare LLM and REINVENT molecule-level distributions within each round using two-sided Mann–Whitney U tests, with Holm correction across the five rounds separately for each organization and metric. n.s., adjusted *p* ≥ 0.05; ∗ for *p* < 0.05; ∗∗ for *p* < 0.01; ∗∗∗ for *p* < 0.001.

**Figure S5:**
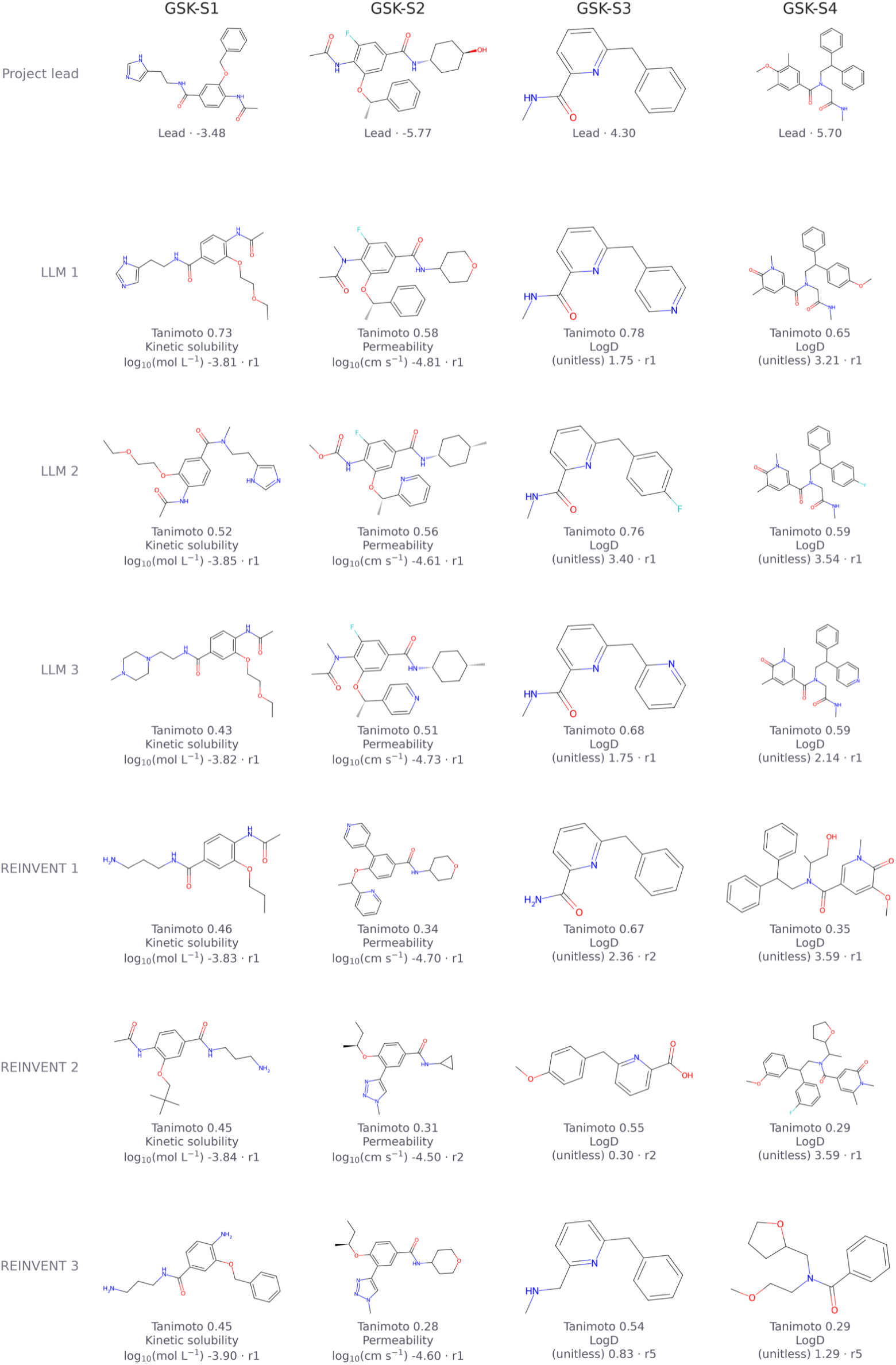
Best held-out compounds and top-ranked LLM and REINVENT proposals for the GSK projects. Best measured held-out molecule plus the top three LLM and top three REINVENT proposals per project.

**Figure S6:**
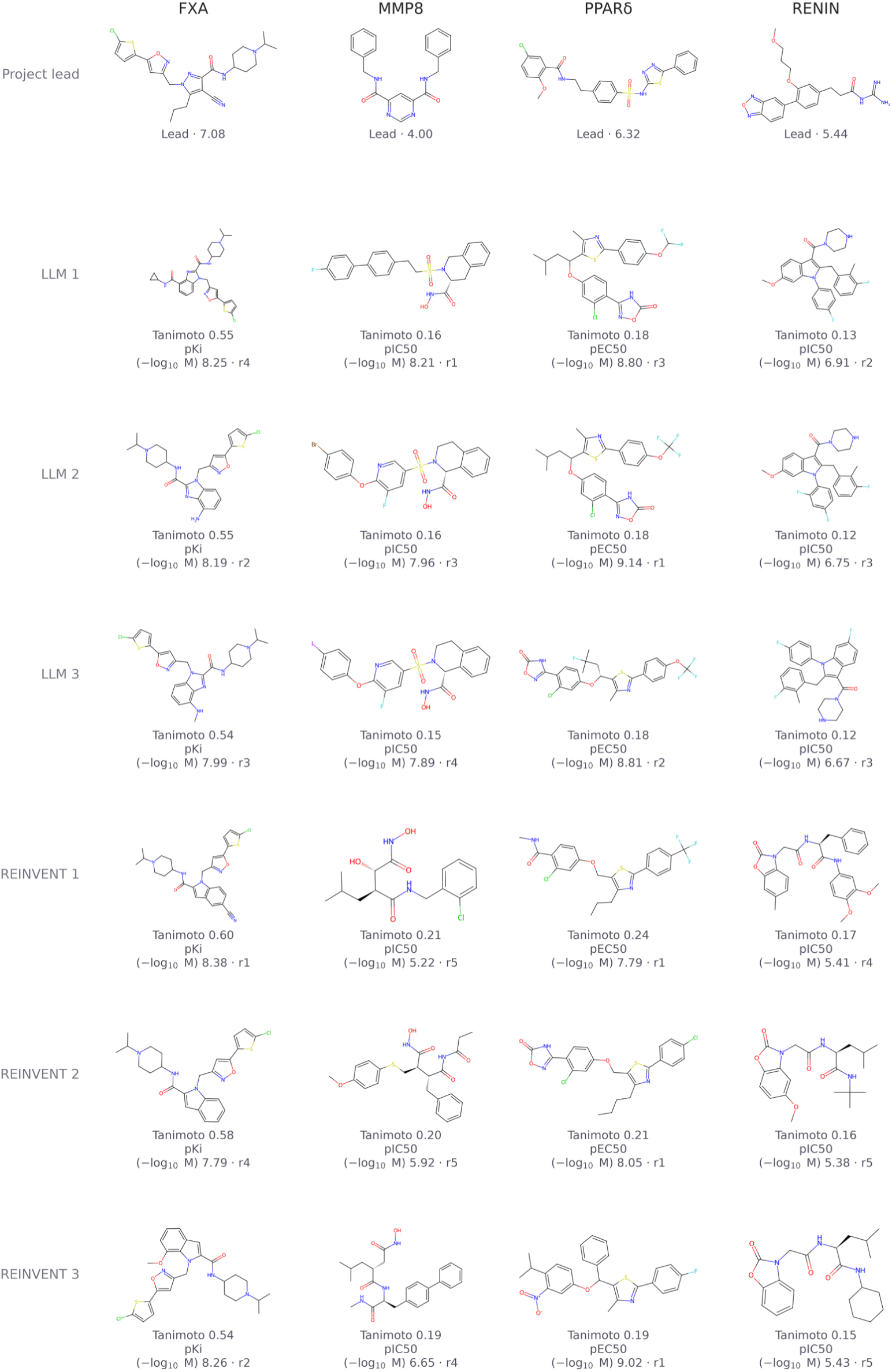
Best held-out compounds and top-ranked LLM and REINVENT proposals for the Sanofi projects. Best measured held-out molecule plus the top three LLM and top three REINVENT proposals per target.

**Figure S7:**
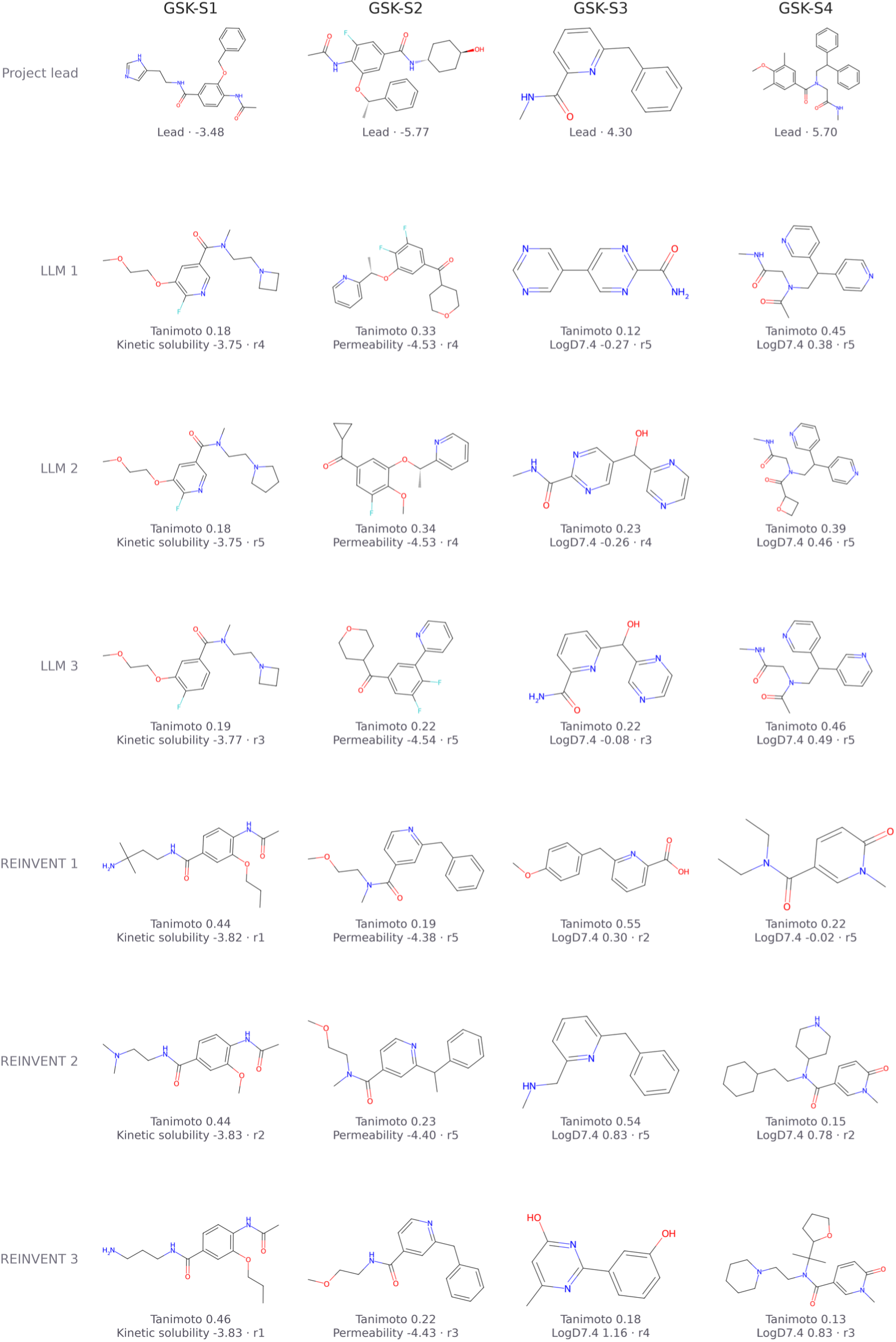
Top five LLM and REINVENT proposals for each GSK project. Columns correspond to GSK-S1–S4; rows show ranks 1–5 for LLM followed by ranks 1–5 for REINVENT. Structures are ranked in the favourable direction of the model-predicted primary endpoint across rounds 1–5 after canonical-SMILES deduplication. The predicted endpoint value and originating round are below each molecule.

**Figure S8:**
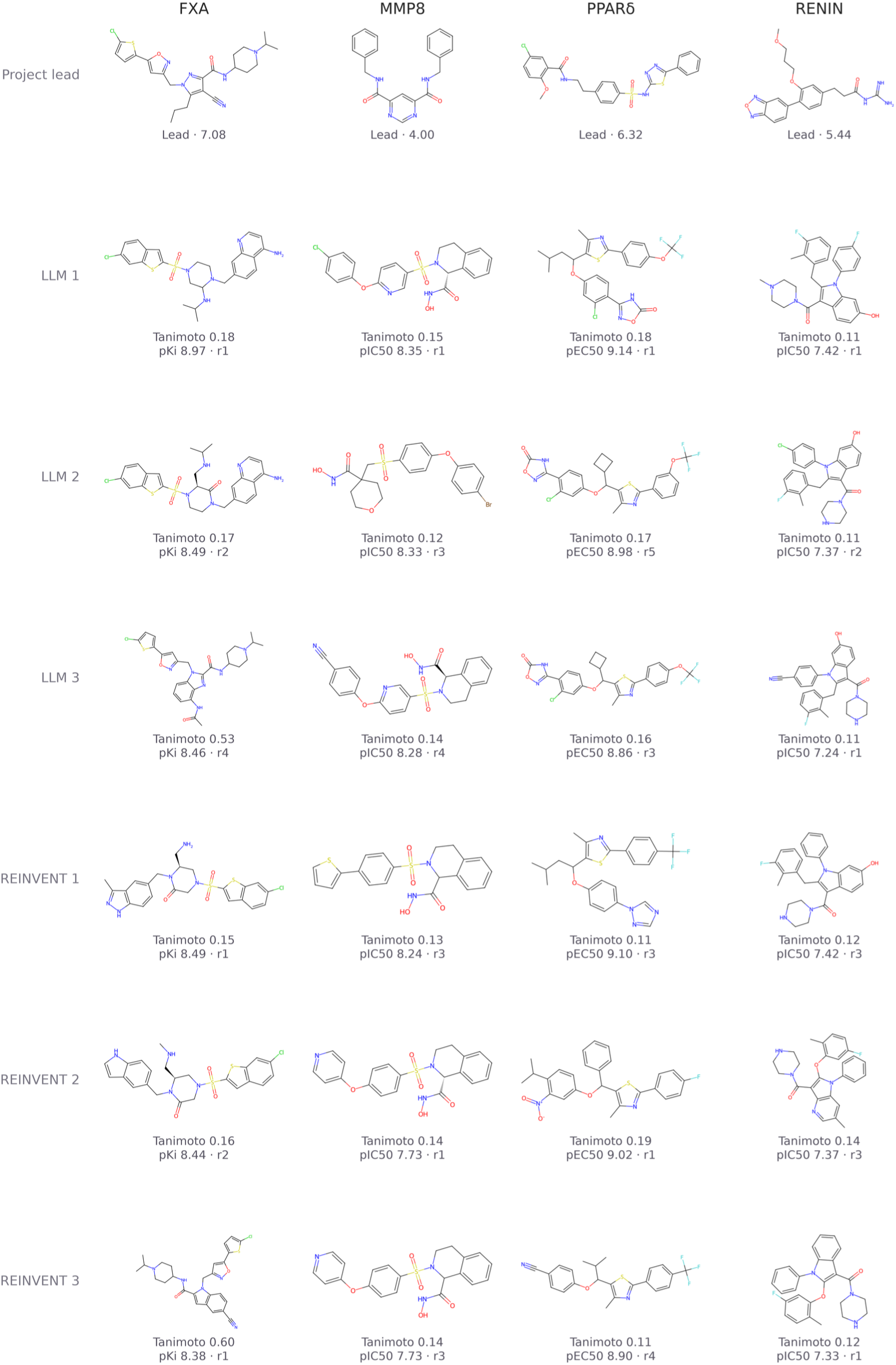
Top five LLM and REINVENT proposals for each Sanofi project. Columns correspond to FXA, MMP8, PPAR*δ* and RENIN; rows show ranks 1–5 for each route. Ranking uses the project model for p*K*_i_, pIC_50_ or pEC_50_ prediction in the favorable direction across rounds 1–5 after canonical-SMILES deduplication. Values and originating rounds are printed below each structure.

**Figure S9:**
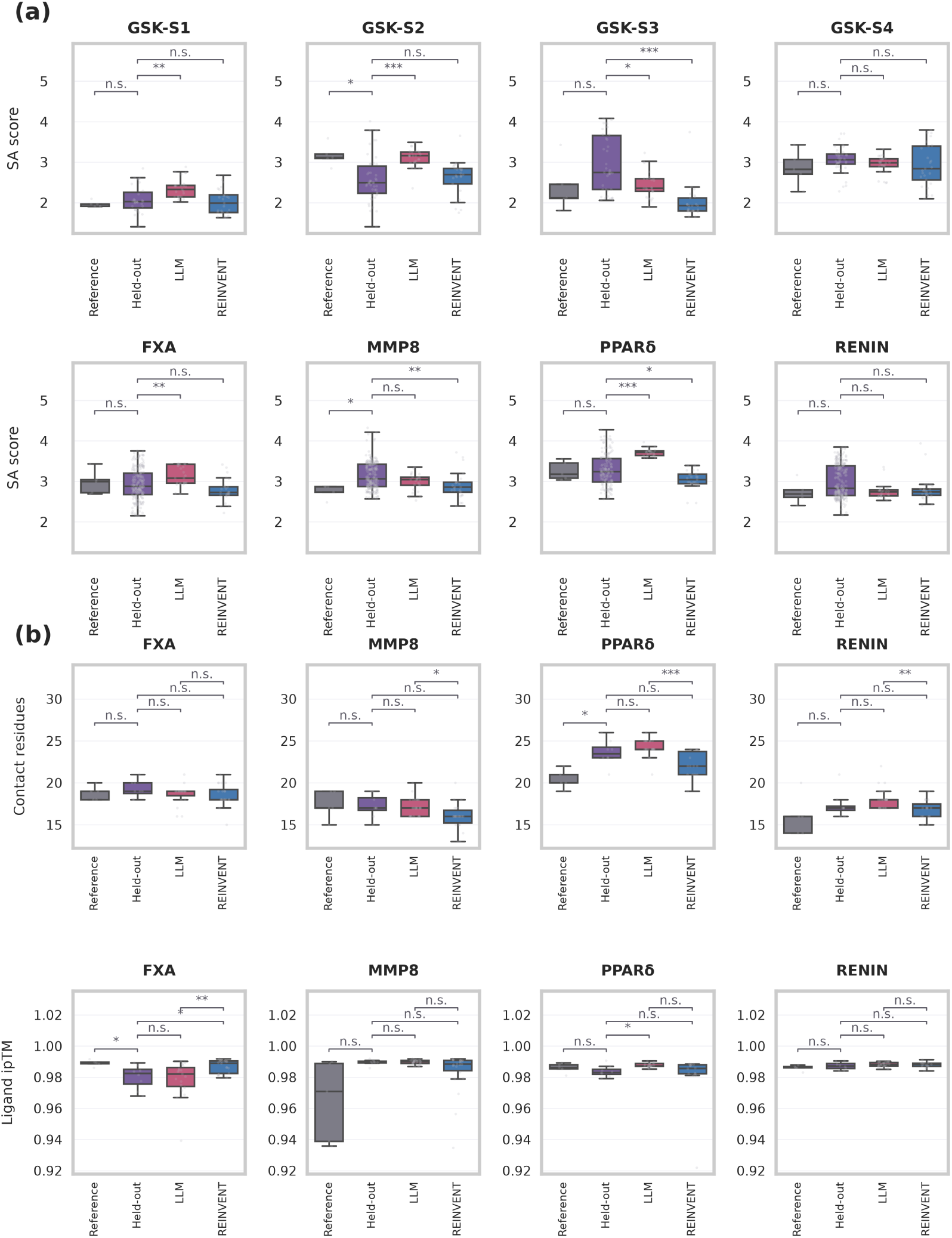
Structural quality of the reference held out and proposed compounds. (a) Synthetic accessibility (SA) scores across the eight projects. Gray, green, red, and blue denote Reference, Held-out, LLM and REINVENT molecules, respectively. (b) Structural quality of predicted complexes. Rows report the number of protein residues with a heavy atom within 4 Å of the ligand and ligand ipTM. Boxes show medians and interquartile ranges; whiskers extend to 1.5 times the interquartile ranges. Brackets compare Held-out–LLM, Held-out–REINVENT and LLM–REINVENT distributions using two-sided Mann–Whitney U tests. n.s., adjusted *p* ≥ 0.05; ∗ for *p* < 0.05; ∗∗ for *p* < 0.01; ∗ ∗ ∗ for *p* < 0.001.

**Figure S10:**
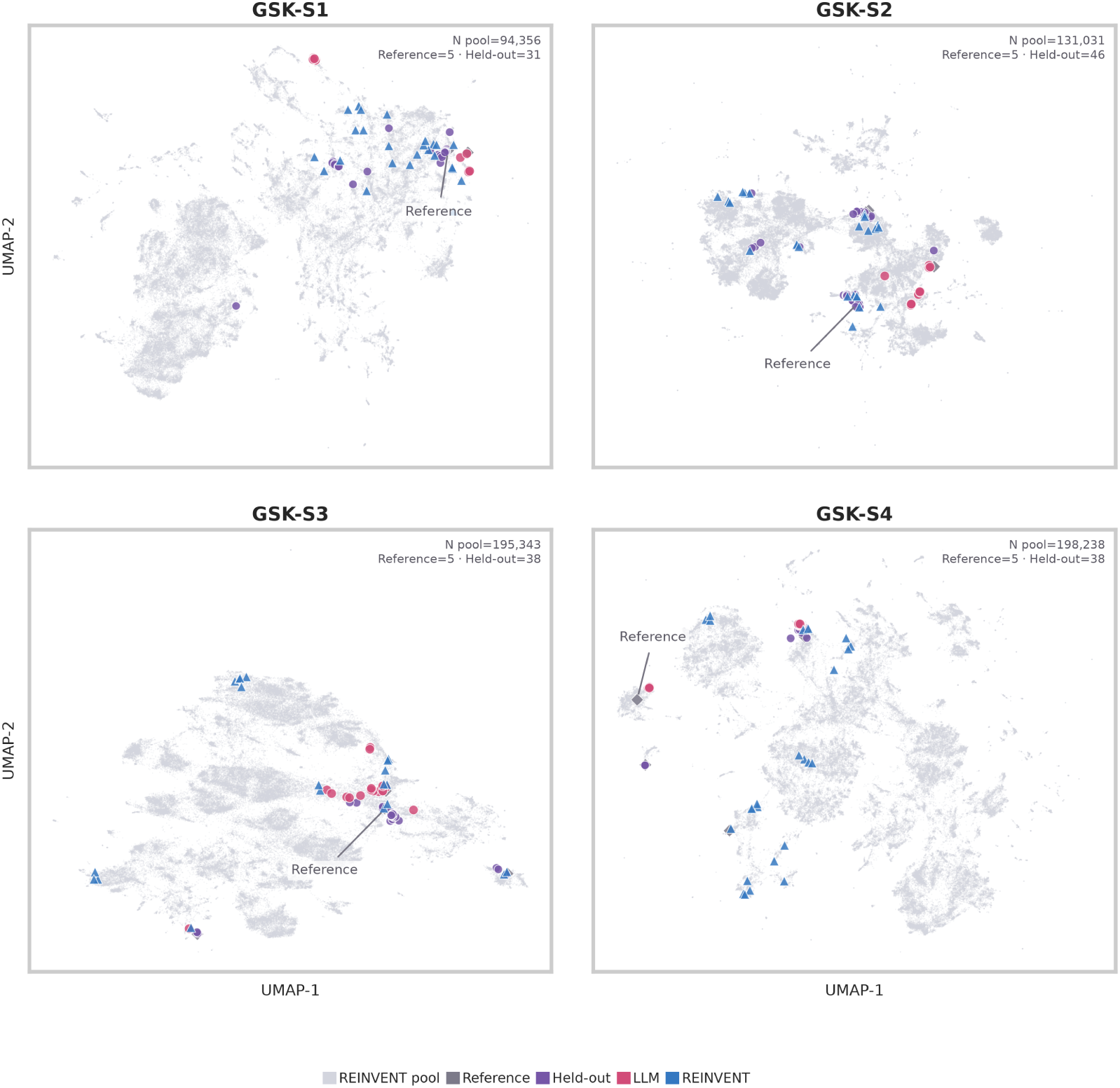
UMAP chemical space coverage for the four GSK projects. Light-gray circles represent unique structures from the internal REINVENT pools, purple diamonds the five project Reference compounds, green circles Held-out compounds, red circles final LLM proposals and blue triangles final REINVENT proposals. Embeddings use 2,048-bit ECFP4 fingerprints and Jaccard distance. Each project is embedded independently. UMAP was computed with umap-learn [63].

**Figure S11:**
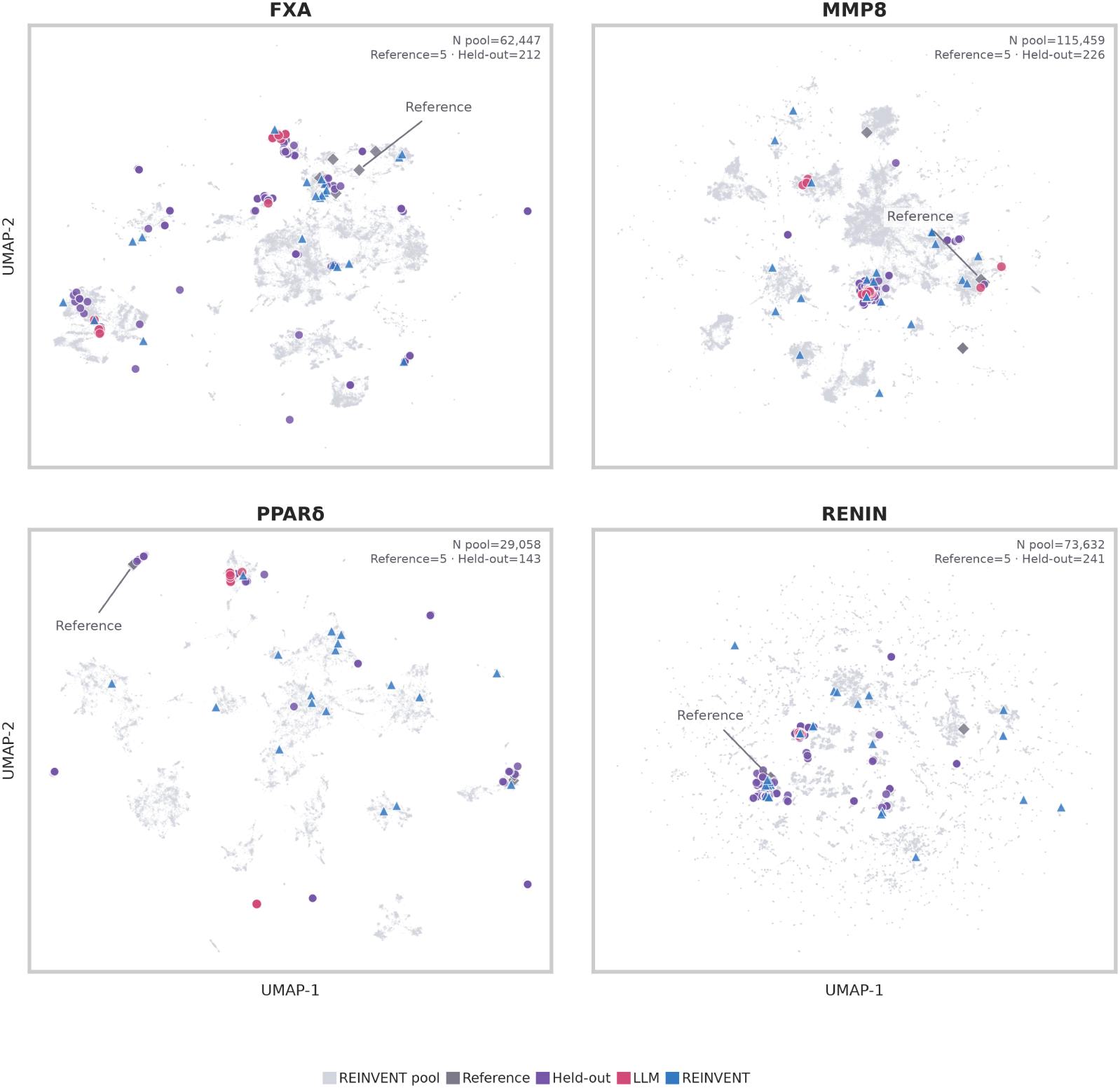
UMAP chemical space coverage for the four Sanofi projects. Light-gray circles represent unique structures from the internal REINVENT pools, purple diamonds the five project Reference compounds, green circles Held-out compounds, red circles final LLM proposals and blue triangles final REINVENT proposals. Embeddings use 2,048-bit ECFP4 fingerprints and Jaccard distance. Each project is embedded independently. UMAP was computed with umap-learn [63].

**Figure S12:**
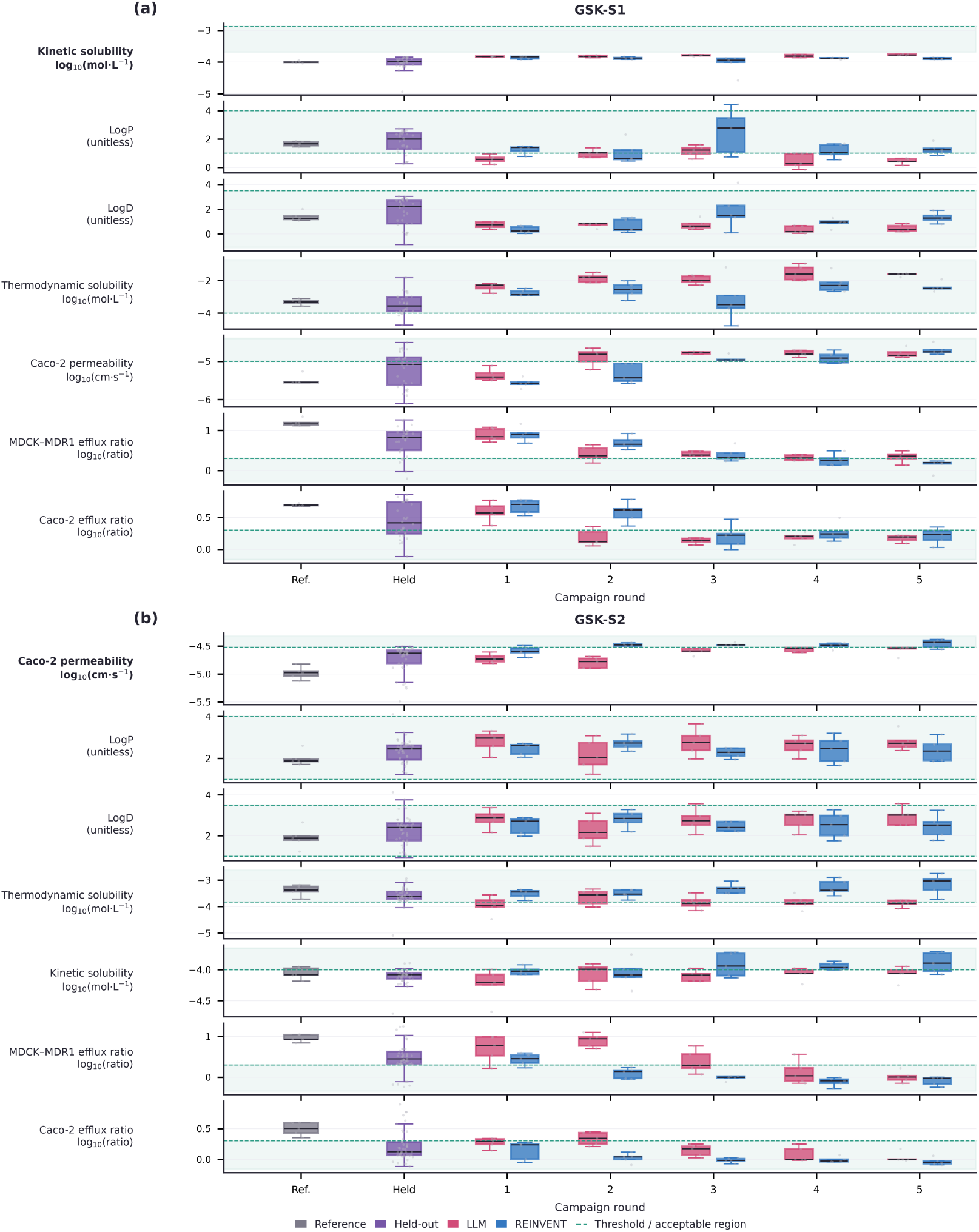
Property distributions during rounds 1–5 of the GSK-S1 and GSK-S2 campaigns. Gray denotes the five shown Reference compounds, green for the published Held-out compounds, red LLM proposals and blue final agent-selected REINVENT proposals. Points represent individual molecules; boxes show the median and interquartile range, with whiskers extending 1.5 times the interquartile range. Each row identifies the endpoint and unit. Dashed green lines mark project-specific criteria where available and generic ADMET context ranges otherwise; pale-green shading shows the corresponding favorable region. Reference and Held-out values are study measurements, whereas proposal values are model predictions.

**Figure S13:**
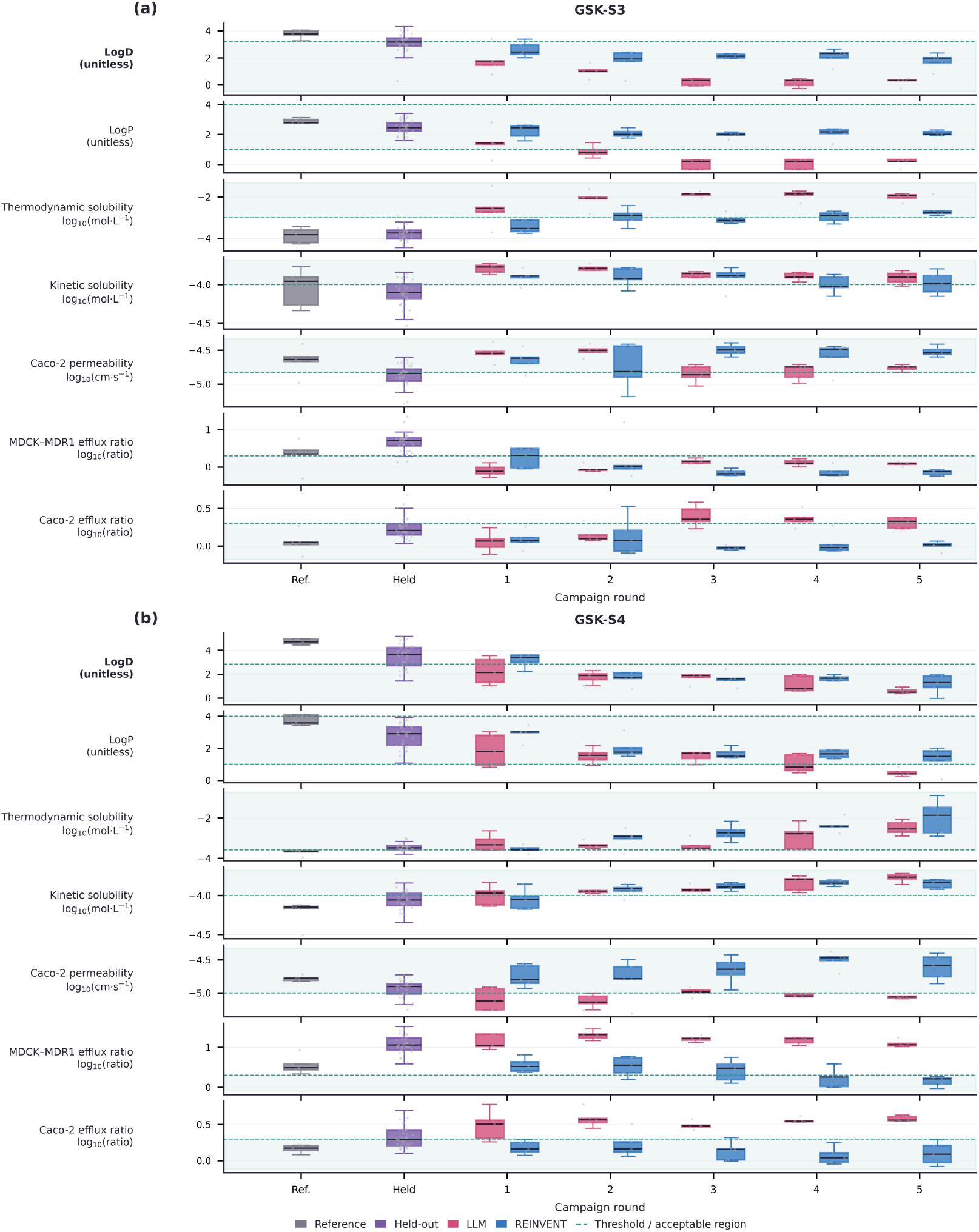
Property distributions during rounds 1–5 of the GSK-S3 and GSK-S4 campaigns. Gray denotes the five shown Reference compounds, green for the published Held-out compounds, red LLM proposals and blue final agent-selected REINVENT proposals. Points represent individual molecules; boxes show the median and interquartile range, with whiskers extending 1.5 times the interquartile range. Each row identifies the endpoint and unit. Dashed green lines mark project-specific criteria where available and generic ADMET context ranges otherwise; pale-green shading shows the corresponding favorable region. Reference and Held-out values are study measurements, whereas proposal values are model predictions.

**Figure S14:**
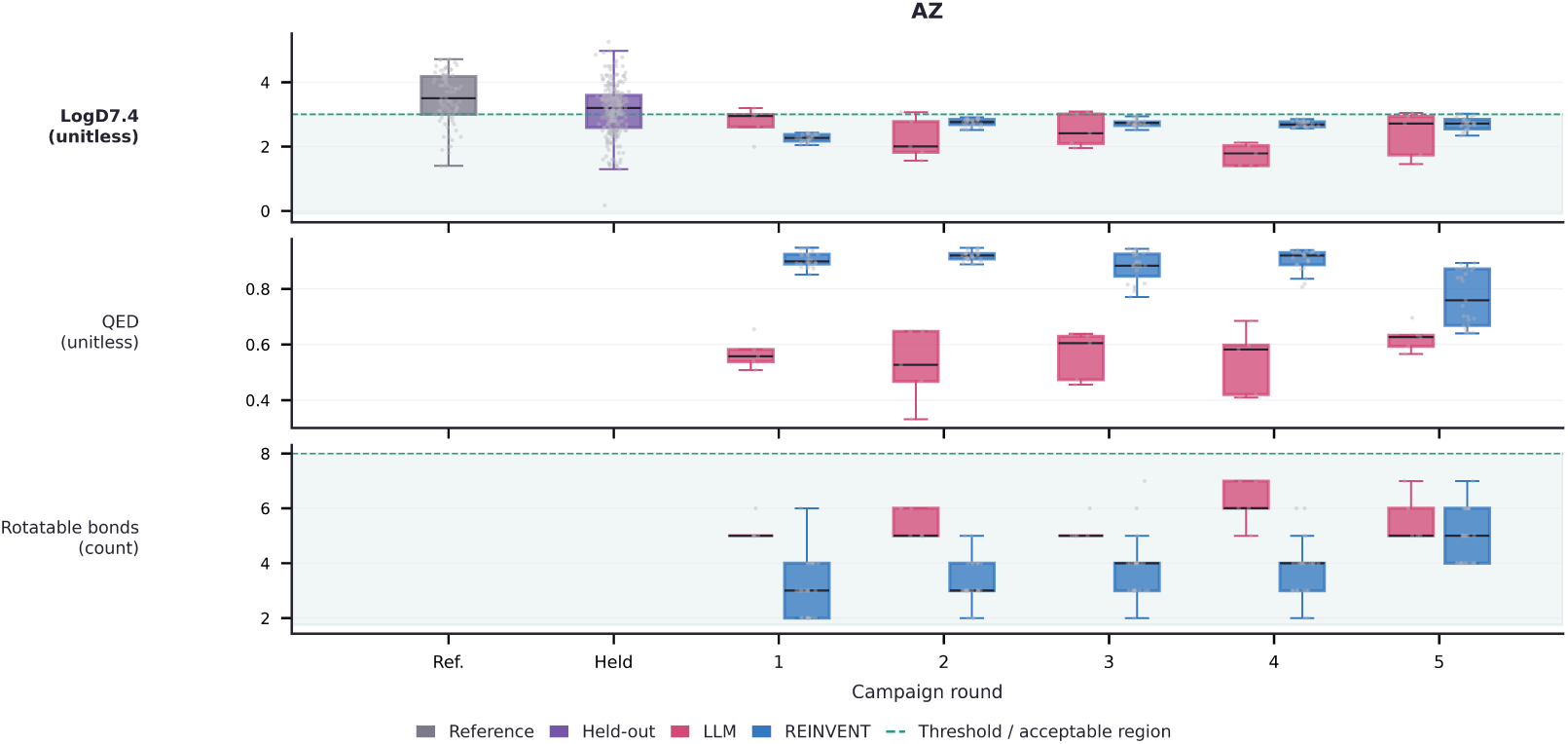
Property distributions during rounds 1–5 of the AstraZeneca log *D*_7.4_ campaign. Gray denotes the five shown Reference compounds, green for the published Held-out compounds, red LLM proposals and blue final agent-selected REINVENT proposals. Points represent individual molecules; boxes show the median and interquartile range, with whiskers extending 1.5 times the interquartile range. Each row identifies the endpoint and unit. Reference and Held-out values are study measurements and are available for log *D*_7.4_ only; QED and rotatable-bond counts are RDKit descriptors computed on the proposals. Dashed green lines mark the campaign criteria, and pale-green shading shows the corresponding favorable region.

**Figure S15:**
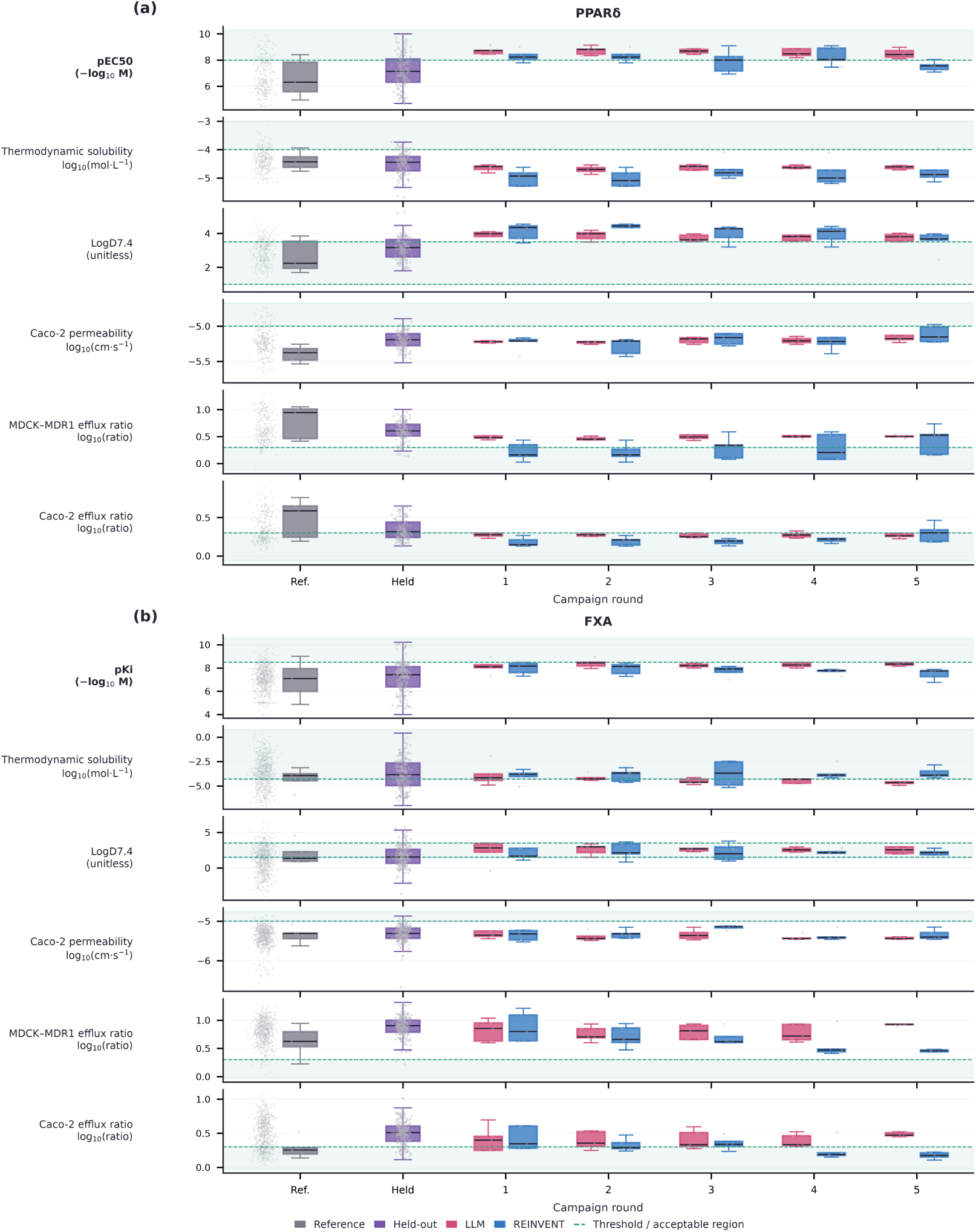
Property distributions during rounds 1–5 of the Sanofi PPARd and FXA campaigns. Gray boxes denote the retrospective five-compound Reference samples used in Figure 1, pale-gray points the complete pre-cutoff project-model training set, green boxes the future Held-out compounds, red boxes LLM proposals and blue boxes final agent-selected REINVENT proposals. The five-compound Reference samples provide visualization context and were not the parents used in the campaigns. Points represent molecules; boxes show medians and interquartile ranges, with 1.5-interquartile-range whiskers. Historical primary-potency values—and FXA solubility and log *D*_7.4_— are experimental measurements. Generic ADMET rows contain model predictions, while TPSA and molecular weight are RDKit descriptors. Dashed green lines show project criteria or generic contextual ranges, and pale-green shading marks the corresponding favorable region.

**Figure S16:**
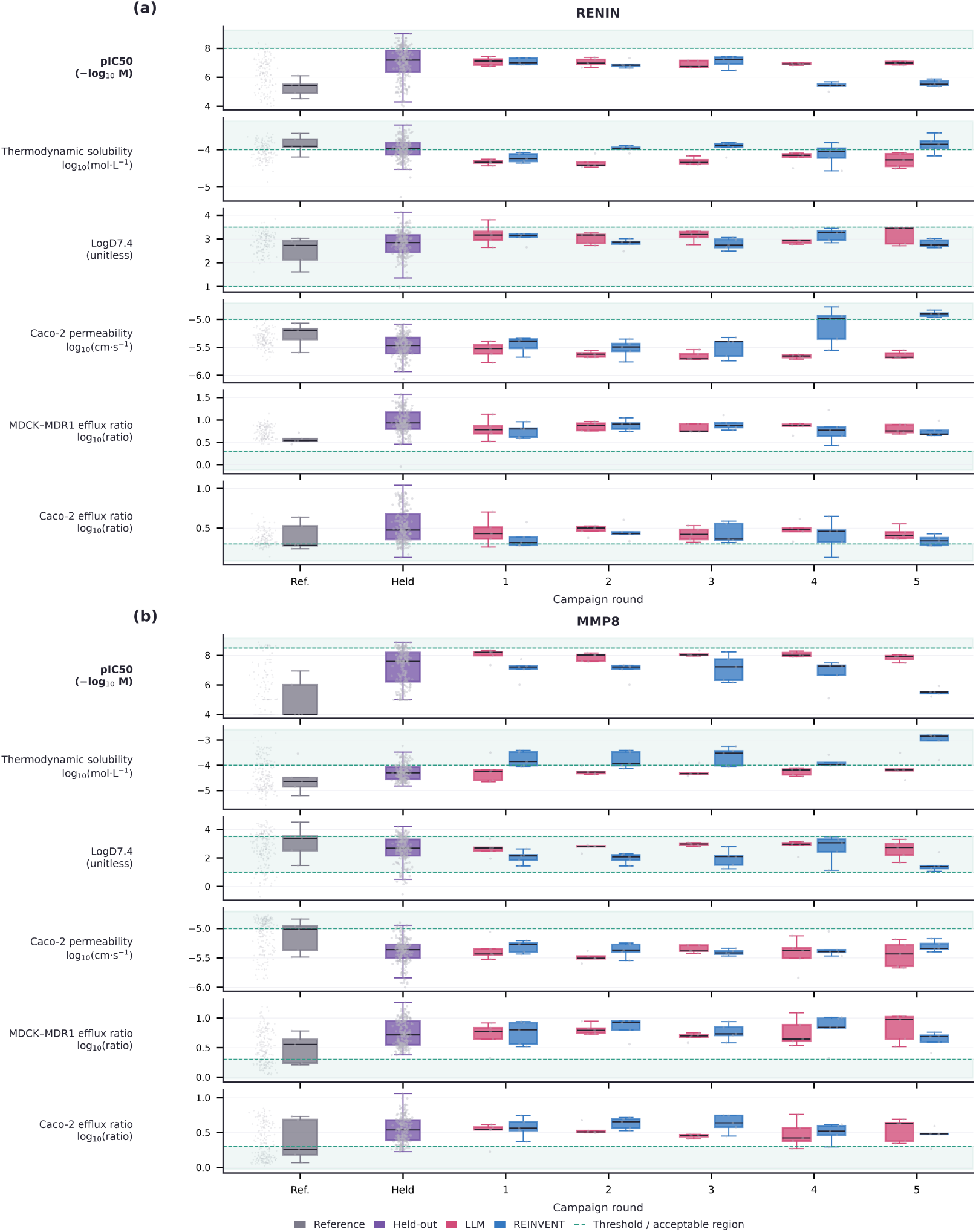
Property distributions during rounds 1–5 of the four Sanofi RENIN and MMP8 campaigns. Gray boxes denote the retrospective five-compound Reference samples used in Figure 1, pale-gray points the complete pre-cutoff project-model training set, green boxes the future Held-out compounds, red boxes LLM proposals and blue boxes final agent-selected REINVENT proposals. The five-compound Reference samples provide visualization context and were not the parents used in the campaigns. Points represent molecules; boxes show medians and interquartile ranges, with 1.5-interquartile-range whiskers. Historical primary-potency values—and FXA solubility and log *D*_7.4_—are experimental measurements. Generic ADMET rows contain model predictions, while TPSA and molecular weight are RDKit descriptors. Dashed green lines show project criteria or generic contextual ranges, and pale-green shading marks the corresponding favorable region.

**Figure S17:**
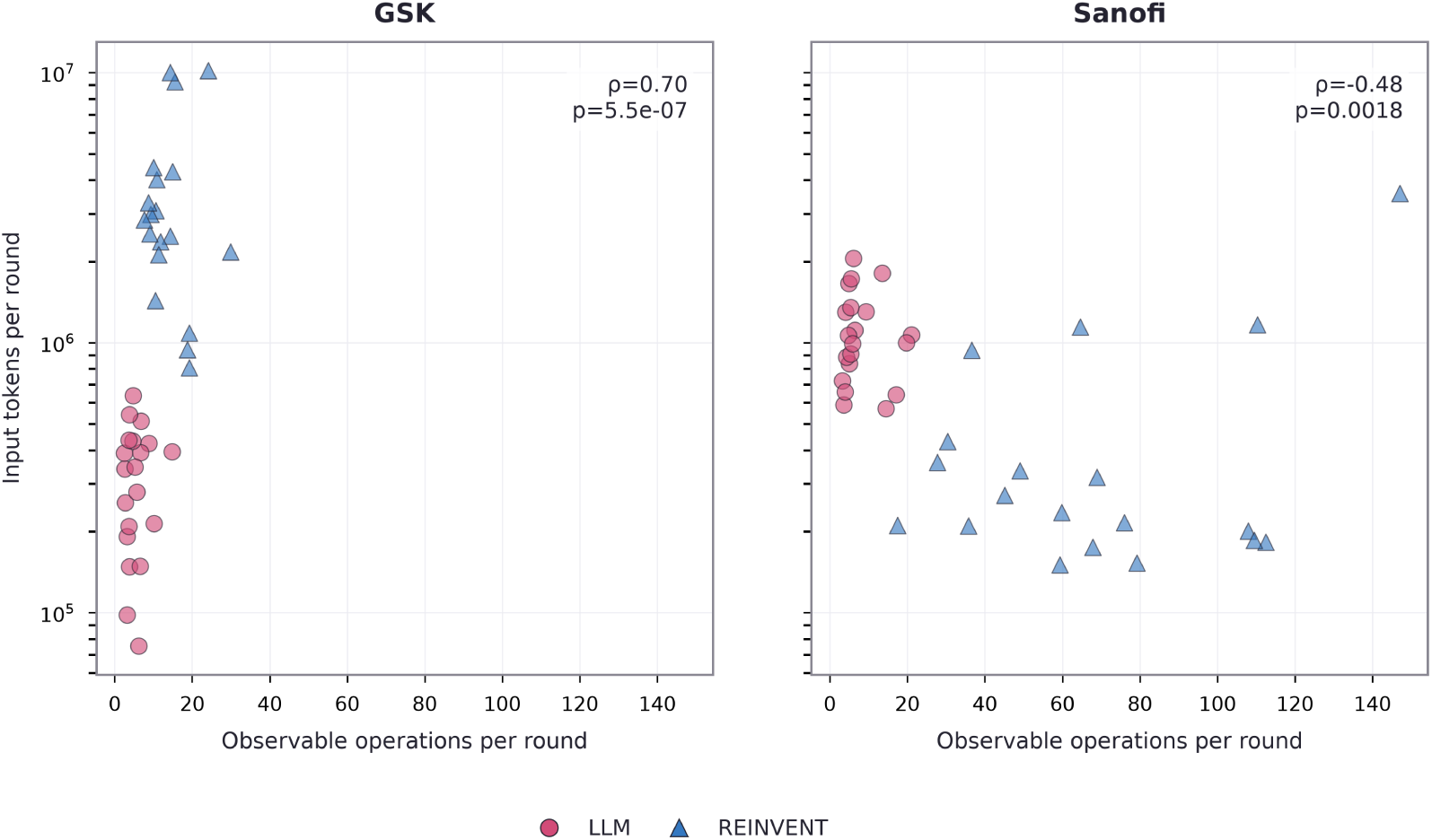
Input-token volume versus observable agent operations during rounds 1–5. LLM and REINVENT values include model-context tokens and agent-visible operations; internal REINVENT scorer calls are excluded.

**Figure S18:**
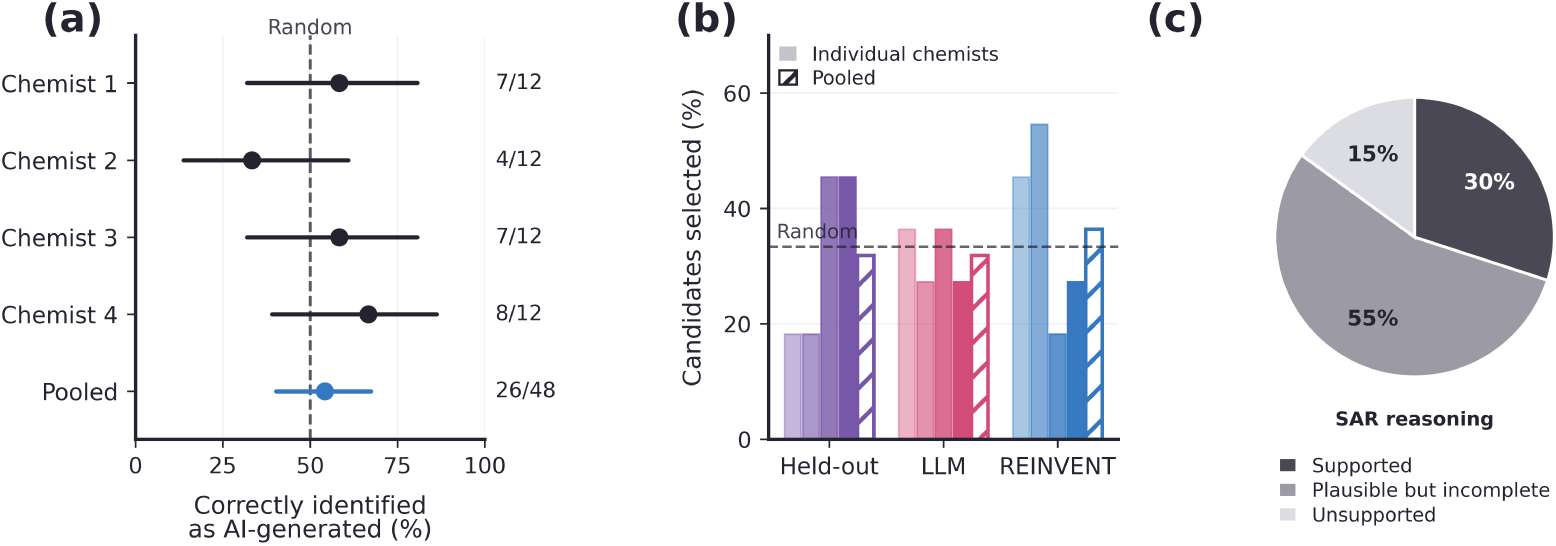
Blinded medicinal-chemist evaluation of agent proposals. Four chemists completed the 28-question instrument described in Section S6.3. **(a) Source discrimination.** For each of the 12 paired cases, the evaluator judged which of two structures was AI-generated; points show the proportion correct with Wilson 95% intervals, and the dashed line marks chance. Counts to the right give correct answers over cases answered. **(b) Candidate prioritization.** Proportion of the 11 prioritization cases in which a held-out, LLM or REINVENT candidate was selected as the next compound to test; solid bars are individual chemists and the hatched bar pools all responses. The dashed line marks the proportion expected if the three sources were chosen at random. **(c) SAR reasoning.** Distribution of verdicts over the 5 rationales, pooled across chemists.

